# Network of hotspot interactions cluster tau amyloid folds

**DOI:** 10.1101/2022.07.01.498342

**Authors:** Vishruth Mullapudi, Jaime Vaquer-Alicea, Vaibhav Bommareddy, Anthony R. Vega, Bryan D. Ryder, Charles L. White, Marc. I. Diamond, Lukasz A. Joachimiak

## Abstract

Cryogenic electron microscopy has revealed unprecedented molecular insight into the conformation of β-sheet-rich protein amyloids linked to neurodegenerative diseases. It remains unknown how a protein can adopt a diversity of folds and form multiple distinct fibrillar structures. Here we develop an *in silico* alanine scan method to estimate the relative energetic contribution of each amino acid in an amyloid assembly. We apply our method to twenty-seven *ex vivo* and *in vitro* fibril structural polymorphs of the microtubule-associated protein tau. We uncover networks of energetically important interactions involving amyloid-forming motifs that stabilize the different fibril folds. We test our predictions in cellular and *in vitro* aggregation assays. Using a machine learning approach, we classify the structures based on residue energetics to identify distinguishing and unifying features. Our energetic profiling suggests that minimal sequence elements that control the stability of tau fibrils, allowing future design of protein sequences that fold into unique structures.

## Introduction

Deposition of β-sheet-rich amyloids are associated with both a diverse group of neurodegenerative and systemic diseases as well as a set of functional amyloid proteins. Amyloidogenic proteins are highly variable in sequence, structure, and cellular function. These proteins assemble into highly stable fibrillar aggregates composed of layers of β-strands oriented orthogonally to the fibril axis [1, 2]. Despite this common “cross-β” architecture and fibrillar morphology, amyloid fibrils adopt a diverse range of structures. In these fibrils, amyloid proteins are folded into monomer layers which are themselves stacked to form an amyloid protofilament. These protofilaments can then associate to form either single or multi-protofilament amyloid fibrils [2].

Aggregation of amyloidogenic proteins is implicated in a variety of diseases, including transmissible spongiform encephalopathies (prion protein/PrP), light chain amyloidosis (immunoglobulin light chain), Parkinson’s Disease (α-synuclein/α-syn), type two diabetes (islet amyloid polypeptide/IAPP), transthyretin amyloidosis (transthyretin/TTR), Alzheimer’s Disease (amyloid-β/Aβ and microtubule-associated protein tau/tau) [3, 4]. Other amyloidoses are similarly related to various forms of systemic or neurological dysfunction. The formation and extension of these fibrils has been theorized to follow prion-like mechanisms, in which a misfolded ‘seed’ protein provides a template that catalyzes the misfolding and fibrillar aggregation of native-state proteins [5].This prion-like misfolding mechanism is implicated in the transmission and propagation of amyloid pathology in the distribution of amyloid fibrils and disease progression within an organism, and (in the case of PrP) between organisms [6, 7]. Prions produce stable, unique conformational ‘strains’, seeds of which reliably propagate both structure and patterns of disease progression. It has been demonstrated that PrP [8], Aβ [9], tau [10] and α-synuclein [11] all adopt conformations capable of propagating aggregate structure and disease pathology, However, until recently, the relation of these strains to amyloid fibril structure has been unclear.

Tau is a key amyloid-forming protein implicated in tauopathies, a group of neurodegenerative disorders that includes Alzheimer’s Disease (AD), Corticobasal Degeneration (CBD), Progressive Supranuclear Palsy (PSP), and others [12]. These tauopathies are characterized by the pathologic aggregation of tau into insoluble amyloid fibrils, and the presence and abundance of these fibrillar deposits are associated with dementia and reduced cognitive function [13]. Under normal conditions, tau is intrinsically disordered and soluble. Tau functions to stabilize microtubules in an extended conformation recently demonstrated in a cryo-EM structure [14]. Over the last five years, advances in cryo-electron microscopy (cryo-EM) and the development of helical reconstruction methods have enabled the structure determination of a variety of fibrillar conformers of tau, each associated with a particular tauopathy [15–22]. Beyond tau, several amyloid proteins including PrP [23], Aβ [24–26], IAPP [27, 28], and α-syn [29–33] have also been shown to adopt multiple structurally distinct conformations, both with and without mutations. These structures have revealed unprecedented molecular details of fibrillar assemblies derived from patient and recombinant sources of tau, α-syn, TTR, IAPP, Aβ, and other proteins, but the molecular rules for how these structures are formed remain poorly understood, particularly *in vivo* [1, 19, 34–36]. At the time of publication there have been nine distinct conformations of tau fibrils extracted from human brain tissue (sixteen, if different protofilament arrangements are included) thus far determined using cryo-EM, highlighting the capacity of a single protein sequence to adopt a diversity of structures associated with distinct disease phenotypes [15–22].

Both the physiological and biophysical determinants of structural polymorphism, as well as the possible range of polymorphic aggregate conformations that a single protein can adopt are uncertain. The mechanisms by which tau adopts aggregation-prone conformations remain unclear, but it has been recently proposed that tau can adopt monomeric aggregation-prone ‘seed’ conformations by rearranging local conformations within the microtubule binding region further stabilized by the termini [37, 38]. Furthermore, it has been proposed that pathologic monomers derived from different tauopathies may encode information to replicate the disease conformation [39] and that these species precede the appearance of soluble oligomers and fibrils [40]. The local motifs within tau that are drivers of aggregation have been proposed to be the two core amyloid-forming elements ^275^VQIINK^280^ and ^306^VQIVYK^311^ located at the beginning of repeat domains two and three, respectively [37]. Additionally, it has been proposed that these amyloid motifs are normally engaged in local interactions within tau to limit their aggregation propensity [41]. Derived from this work, it has been proposed that local structural rearrangements surrounding the amyloid motifs encoded in a tau monomer may drive differentiation into distinct structural polymorphs. Thus, it is possible that specific misfolding events in a monomer are sufficient to initiate assembly of tau into conformationally distinct aggregates.

To explore the determinants of amyloid polymorphism, we use tau as a model protein to understand how different fibril folds may form and what interactions may mediate their stability. We note that amyloidogenic motifs in tau play important roles in stabilizing heterotypic nonpolar contacts within tau fibrils. To further understand the interactions responsible for stabilizing amyloid fibrils, we deploy an *in silico* method using Rosetta to probe residue energetics in across different fibrillar structures. We first develop a minimization protocol for fibrils yielding minimized structures that retain near native backbone conformations and recapitulate sidechain rotamers and interactions. Using this platform, we implement an alanine mutagenesis scan for twenty-seven *ex vivo* and recombinant tau fibril structures and estimate the relative energetic contribution of specific residues to the stability of tau fibrils. We use these estimated contributions to identify the thermodynamic hotspots that contribute to and differentiate the multiplicity of fibril polymorphs found in tauopathies. We uncover key hotspot residues involving amyloidogenic motifs and identify a modular network of interactions of which subsets are preserved across structurally diverse folds. We then test the role of these heterotypic interactions in peptide co-aggregation and in cell experiments and show that amyloid-motif aggregation can be regulated by heterotypic contacts. Finally, we use a machine learning approach to uncover key energetic features that will help classify the different structures.

## Results

### Tauopathy fibril cores involve modular interactions with amyloid motifs

To understand how primary sequence properties may underpin tau’s amyloid polymorphism, we compared tau’s polar and nonpolar amino acid distribution with its predicted aggregation propensity. Aggregation-promoting regions (APRs) of tau have previously been identified, and prior work suggests sequences surrounding these APRs may regulate their contribution to tau self-assembly [41, 42]. Structure-based computational methods can efficiently predict aggregation-promoting elements (e.g., AmylPred, ZipperDB, Waltz) and have uncovered key motifs including ^275^VQIINK^280^ and ^306^VQIVYK^311^ as APRs central to tau aggregation, but other sequences that are predicted to be weakly amyloidogenic retain increased hydropathy (Fig. 1a) and may be involved in hetero-typic interactions with core amyloid motifs [43–47]. Indeed, ^275^VQIINK^280^ and ^306^VQIVYK^311^ aggregate into ordered fibrils but other sequences predicted to be close to the energetic threshold (Fig. 1a, dashed red line) predicted in ZipperDB form a distribution of ordered aggregates and disordered (i.e., amorphous) aggregates (Fig. 1b). Sequences close to or below this threshold remain soluble (Supplementary Fig. 1a) and do not yield Thioflavin T (ThT) fluorescence signal, a reporter of ordered β-sheet structure formation (Supplementary Fig. 1b). Focusing on the well-characterized ^306^VQIVYK^311^ peptide, we wanted to understand which residues in this motif are important for fibril formation. To test this directly, we measured the aggregation capacity of ^306^VQIVYK^311^ peptide mutants substituted with alanine at each position in a ThT fluorescence aggregation assay. We find that alanine mutations at positions one, three, and four (i.e., **A**QIVYK, VQ**A**VYK and VQI**A**YK) completely abolish ThT fluorescent signal while substitutions at positions two and six (i.e., V**A**IVYK and VQIVY**A**) retain near WT aggregation properties (Supplementary Fig. 1c). Position five (i.e., VQIV**A**K) gives intermediate fluorescence signal (Supplementary Fig. 1c), but this may be due either to the loss of the aromatic residue which is important for ThT binding or to a decrease in fibrilization [48]. We confirmed our ThT results by imaging each sample using Transmission Electron Microscopy (TEM) and find that the peptides that yield positive ThT signal have fibrils. One exception is the **A**QIVYK peptide, which yielded thin fibrils that were negative in the ThT assay suggesting that this mutation alters the morphology of the aggregates compared to the other ThT positive structures (Supplementary Fig. 1d). To interpret these data in the context of existing structures of ^306^VQIVYK^311^ and related sequences determined by X-ray crystallography, we find that indeed valine and isoleucine at positions 306 (i.e., **V**QIVYK) and 308 (i.e., VQ**I**VYK) of VQIVYK, respectively, appear to be central in stabilizing intermolecular contacts in either a head-on or offset register (Supplementary Fig. 1e). Additionally, valine at position 309 (i.e., VQI**V**YK) is important despite not forming the central hydrophobic core indicating that additional secondary interactions on both hydrophobic sides of the β-sheet are important for fibril formation. These data suggests that ^306^VQIVYK^311^ has the capacity to stabilize self-interactions in different arrangements in simple peptides using both sides of the β-sheet. Unsurprisingly, if we compare the placement of ^306^VQIVYK^311^ in the nine unique *ex vivo* tau protofilament polymorphs determined to date (interpreting only a single layer), we observe that ^306^VQIVYK^311^ is engaged in interactions with different sequences from repeat domains two and four (Supplementary Fig. 2a) that are often aggregation-prone and match the nonpolar composition of this amyloid motif (Supplementary Fig. 2b, c) [15, 17, 18, 20, 21].

**Figure 1.**
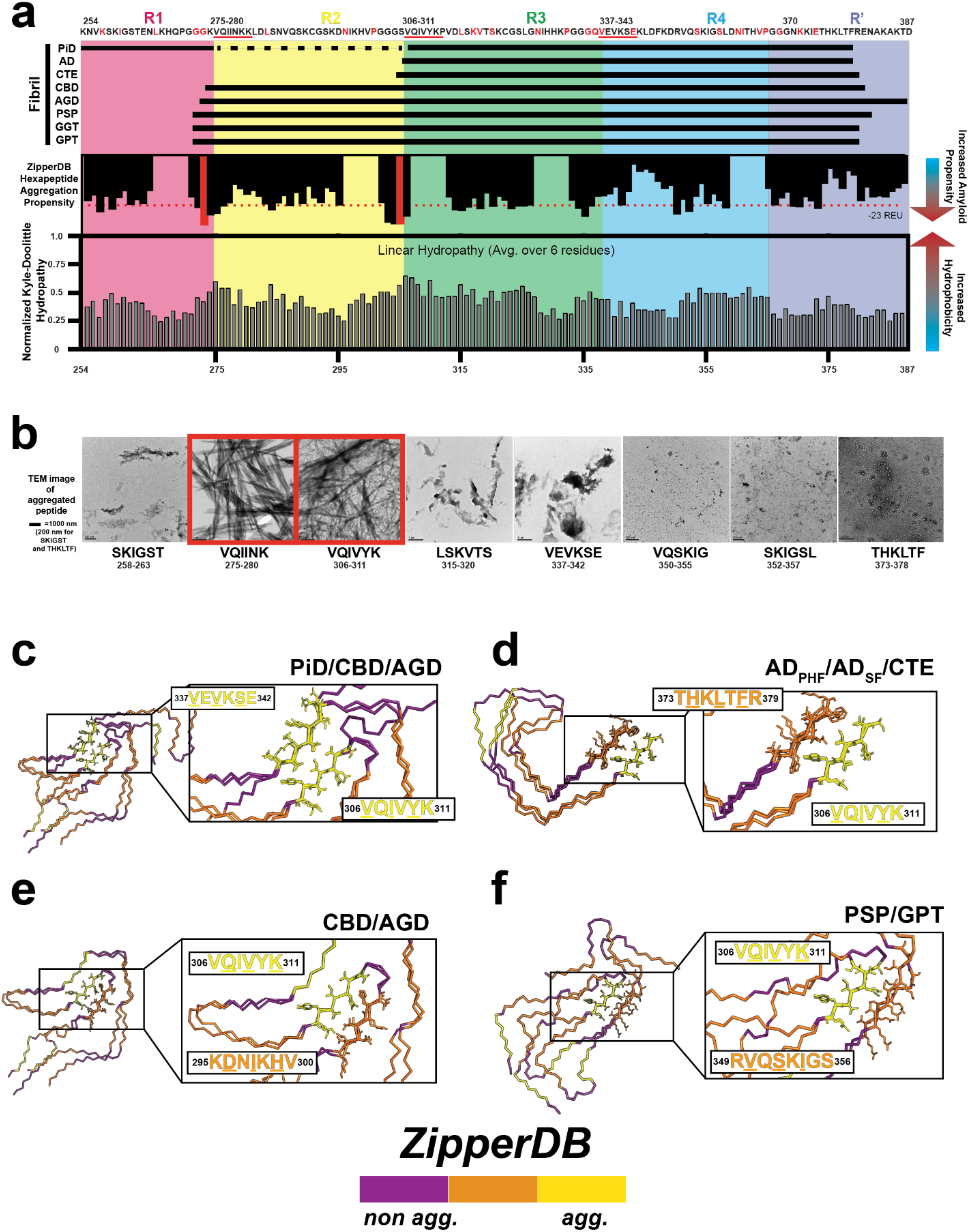
Nonpolar and amyloidogenic core fragments are modular in tau fibrils. **(a)** Diagram of the tau fragments utilized in the fibrillar assemblies highlighting the aggregation propensity of the tau sequence (as estimated by ZipperDB) and its hydrophobicity (shown as Kyte-Doolittle hydropathy, calculated by localCider) [46, 47, 66]. The fragments for each structural polymorph are shown as horizontal bars. The tau repeat domains are colored red, yellow, green, blue and navy blue for R1, R2, R3, R4 and R’. The corresponding tau sequence is shown on the top. Known mutations are colored in red and derived from Alzforum [67]. Key amyloid motifs are underlined. Aggregation propensity is shown as black bars. Dotted red line denotes threshold for high aggregation propensity. The two most highly aggregation-prone sequences, ^275^VQIINK^280^ and ^306^VQIVYK^311^ are highlighted in red. Hydropathy plot is shown as grey bars. **(b**) TEM images of predicted amyloid motifs. ^306^VQIVYK^311^ and ^275^VQIINK^280^ are highly aggregation prone (boxed in red), and form ordered assemblies while the other sequences predicted to have lower aggregation propensity form disordered or amorphous aggregates. **(c-f**) Comparison of ^306^VQIVYK^311^ amyloid motif interactions across structural polymorphs. **(c**) ^306^VQIVYK^311^ forms conserved nonpolar interactions with ^337^VEVKSE^342^ in PiD, CBD and AGD structures. **(d**) ^306^VQIVYK^311^ forms conserved nonpolar interactions with ^373^THKLTFR^379^ in AD-PHF, AD-SF and CTE structures. **(e)** ^306^VQIVYK^311^ forms nonpolar interactions with ^294^KDNIKHV^300^ in AGD and CBD structures. **(f)** ^306^VQIVYK^311^ forms nonpolar interactions with ^349^RVQSKIGS^356^ in PSP and GPT structures. The structures are shown as a single layer in a ribbon representation and are colored by amyloidogenic propensity: yellow (high), orange (medium) and purple (low). Amino acid sequences of interacting motifs are colored by aggregation propensity and amino acids important for interaction are underlined.

To look at specific conservation of the interactions with ^306^VQIVYK^311^ we aligned a layer from each of the nine prototypical *ex vivo*-derived structural polymorphs to the ^306^VQIVYK^311^ amyloid motif. We find that ^306^VQIVYK^311^ makes conserved interactions to other predicted amyloid motifs in subsets of the fibrillar structures (Fig. 1c-f). In the PiD, CBD and AGD structures, V306, I308 and Y310 from VQIVYK makes identical interactions to V337 and V339 from ^337^VEVKSE^342^ (Fig. 1c). In the simpler, AD-PHF, AD-SF and CTE folds, V306, I308 and Y310 from ^306^VQIVYK^311^ again interacts with L376 and F378 from ^373^THKLTFR^379^ (Fig. 1d). In the CBD and AGD folds, V306, I308, Y310 and K311 from ^306^VQIVYK^311^ interacts with D295, I297 and H299 of ^295^DNIKHV^300^ highlighted by nonpolar contacts between V306 and I297 but also a buried salt bridge between K311 and D295 (Fig. 1e). Interestingly, in GGT I297 from ^295^DNIKHV^300^ interacts with ^306^VQIVYK^311^ but the other residues are out of register. Finally, in the PSP and GPT structures, Q307, V309, and K311 of ^306^VQIVYK^311^ interact with V350, S352 and I354 of ^349^RVQSKIGS^355^ (Fig. 1f). These analyses suggest that aggregation-prone elements including ^306^VQIVYK^311^ are used modularly to bury nonpolar contacts in the cores of *ex vivo*-derived tau fibril structures. Our data indicate that the interactions of key amyloid sequences with other hydrophobic sequence elements play important roles in tau amyloid assembly and the diversity of these possible stabilizing interactions may be central to the formation of a diversity of structural polymorphs. Importantly, we predict that ^306^VQIVYK^311^ is a key regulator of tau assembly which uses one or two surfaces to stabilize hydrophobic interactions in simple and complex fibril cores. The fibrils can be classified into two general categories: one where the ^306^VQIVYK^311^ peptide strand interacts with a second β-strand in a one-sided β-sheet interaction, and another in which ^306^VQIVYK^311^ engages two other β-strands in a two-sided β-sheet interaction. We classify CBD, AGD, PSP, GGT and GPT as fibrils with two-sided ^306^VQIVYK^311^ interactions, while AD, CTE and PiD (and the heparin derived fibril structures) are comprised of monomer layers with one-sided ^306^VQIVYK^311^ interactions.

To gain a more coarse-grained view of the types of residues that are buried in the nine prototypical *ex vivo* structures we colored each structure by amino acid polarity. We find that nonpolar amino acids are often buried in the fibril cores (Supplementary Fig. 2c, yellow spheres) while basic residues are presented on the outside of the fibrils (Supplementary Fig. 2c, blue spheres). We also quantified the degree of burial of different amino acid types by using Rosetta to calculate the change in solvent accessible surface area between a fully extended monomer to its folded monomer conformation alone (ΔSASA_monomer_), or to its folded monomer in the context of a fibril (ΔSASA_fibril_). Aggregating this data over the nine *ex vivo* prototypical fibril polymorphs, we find that nonpolar amino acids have both a large ΔSASA_monomer_ and ΔSASA_fibril_ (Supplementary Fig. 3a, orange), suggesting nonpolar residues are buried when the tau monomer folds into a fibril conformation, and further buried as the additional layers of the fibril form. Polar and acidic residues appear to be distributed more evenly between the core and the surface (Supplementary Fig. 2c) and show similar burial patterns by ΔSASA*^folding^_single layer_* and ΔSASA*^folding^_fibril_* (Supplementary Fig. 3a, green and red). Interestingly, when we quantify change in solvent accessibility for basic residues (mostly lysines) we find that ΔSASA*^folding^_single layer_* is small but ΔSASA*^folding^_fibril_* is large, suggesting that basic residues are solvent exposed but can bury their sidechains in the fibril by stacking their aliphatic sidechains (Supplementary Fig. 2c and 3a, blue). Consistent with this observation a recent study showed that a tau fragment can assemble into AD-PHF and CTE conformations using specific salts indicating that cation and anion interactions with tau may promote tau folding by stabilizing nonpolar burial and screening electrostatic interactions [49]. This analysis indicates that nonpolar residues are generally buried in tau fibril structures-often through interactions with amyloid motifs, likely contributing to the stability of the different protofilament folds.

### Nine layer fibril stack retains native-like properties through minimization and mutagenesis

We developed a framework to interpret the interaction energies across a panel of fibrillar structures. First, we used Rosetta to minimize each fibril structure while retaining the native conformation. Then we used a Flex-ddG based protocol to perform alanine-scanning mutagenesis which allows us to infer the energetic contribution of amino acids in each structure (Fig. 2) [50]. Our protocol starts with generation of coordinate and pairwise-atom structural restraints derived from the starting cryo-EM structures that are employed to minimize the structures. The backrub protocol is implemented to improve sampling of the backbone and sidechain conformational space and improve energy minimization (Fig. 2) [51]. Using these sampled structures, thirty-five wildtype and mutant structures are generated and undergo sidechain optimization and global minimization utilizing the initially developed restraints.

**Figure 2.**
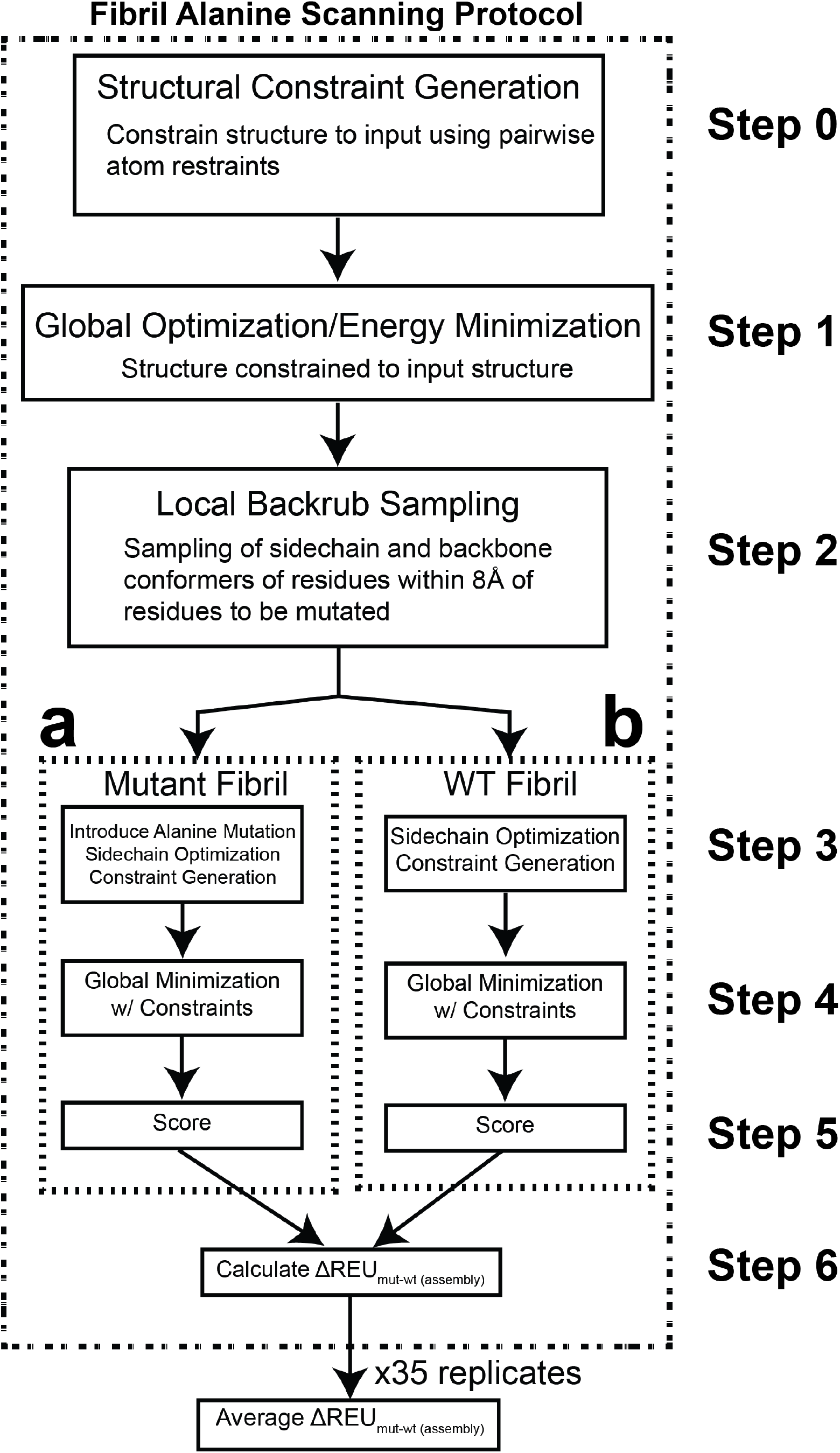
Diagram for the fibril Flex ddG energy estimation method. Outline of fibril flex ΔREU^assembly^_mut-wt_ method steps. This procedure is based on the method described in [50]. (0) The fibril is constrained to the input fibril structure with atom pair constraints with max distance 9.0 Å. (1) Using these constraints, the input fibril structure undergoes energy minimization. Minimizations are performed with constraints that harmonically restrain pairwise atom distance to their values in the input structure. Minimization is run until convergence (absolute score change upon minimization of less than one REU (Rosetta Energy Unit)). (2) The backrub method (the Rosetta BackrubProtocol mover) is used sample additional backbone and sidechain conformers proximal to the mutation site. Each backrub move is undertaken on a randomly chosen protein segment consisting of three to twelve adjacent residues with a C-β atom (C-α for glycines) within 8 Å of the alanine mutant positions, or that residue’s adjacent N and C-terminal residues. All atoms in the backrub segment are rotated locally about an axis defined as the vector between the endpoint C-α atoms. Backrub is run at a temperature of 1.2 kT, for 35000 backrub Monte Carlo trials/steps with default Rosetta settings for backbone angles. 35 ensemble models are generated. (3A) Alanine mutants are introduced to the backrub-sampled model, and sidechain conformations for the mutant structure are optimized using the Rosetta packer. (3B) alongside the alanine mutant model, a wild-type model is also optimized with the packer. (4A) The mutant model is minimized using pairwise atom-atom constraints to the input structure. Minimization is run with the same parameters as in step 1; the coordinate constraints used in this step are taken from the coordinates of the step 3A model. (4B) Same as step 4A, but for the wild-type model. (5A) The model is scored both as a ‘bound’ protein complex and as a split, ‘unbound’ complex. The scores of the split, unbound complex partners are obtained simply by moving the complex halves away from each other. No further minimization or sidechain optimization is performed on the unbound partners before scoring. (5B) similarly to step 5A, the minimized wild-type model is scored as both in a ‘bound’ and ‘unbound’ state (6) The interface ΔΔG and ΔREU^assembly^_mut-wt_ score using the following equations:

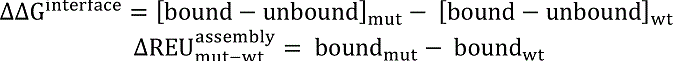

These ΔΔG^interface^ and ΔREU^assembly^_mut-wt_ values are then averaged over the 35 replicates in the ensemble to generate the final ΔΔG and ΔREU^assembly^_mut-wt_ for the residue.

Unlike with globular proteins which can exist stably as monomeric or in a set of defined oligomeric states, fibrils exist as a set of layered monomers without a specific required length/number of layers. To determine a sufficient number of monomer layers for use in the fibril simulations we minimized the AD-PHF and CBD structures using different numbers of monomer layers from a trimer to a nine-mer representing the fibril (Fig. 3a). Minimization yielded energetically favorable assemblies compared to the starting fibril structures with the total energy of the assembly scaling with the number of layers for both wild-type and alanine mutants (Supplementary Fig. 3b, c). Additionally, we computed RMSD distributions for AD-PHF and observe that the RMSD decreases with increasing numbers of layers (Fig. 3b, blue) while for CBD the RMSD remains flat across the range of layers (Fig. 3c, blue) for both wild-type and alanine substituted structures. We suspect the number of layers may help retain the assembly in a native conformation and may help with fibrils with simpler topologies (i.e., improved AD more than CBD). For more in-depth analysis across tau fibrils, we decided to move forward with nine-mer assemblies which balanced low rmsd with cost of computation time. Comparing the minimized nine-mer AD-PHF and CBD structures to the native conformation, we find that our minimization protocol maintains near native backbone and sidechain rotamers in the AD and CBD structures with all-atom RMSDs of 0.486 Å and 0.539 Å, respectively (Fig. 3d, e, blue). Importantly, we recover the native rotamers of sidechains in the core of the fibril, though residues with sidechains on the outside surface that mainly interact with the solvent are more variable (Fig. 3d, e, blue).

**Figure 3.**
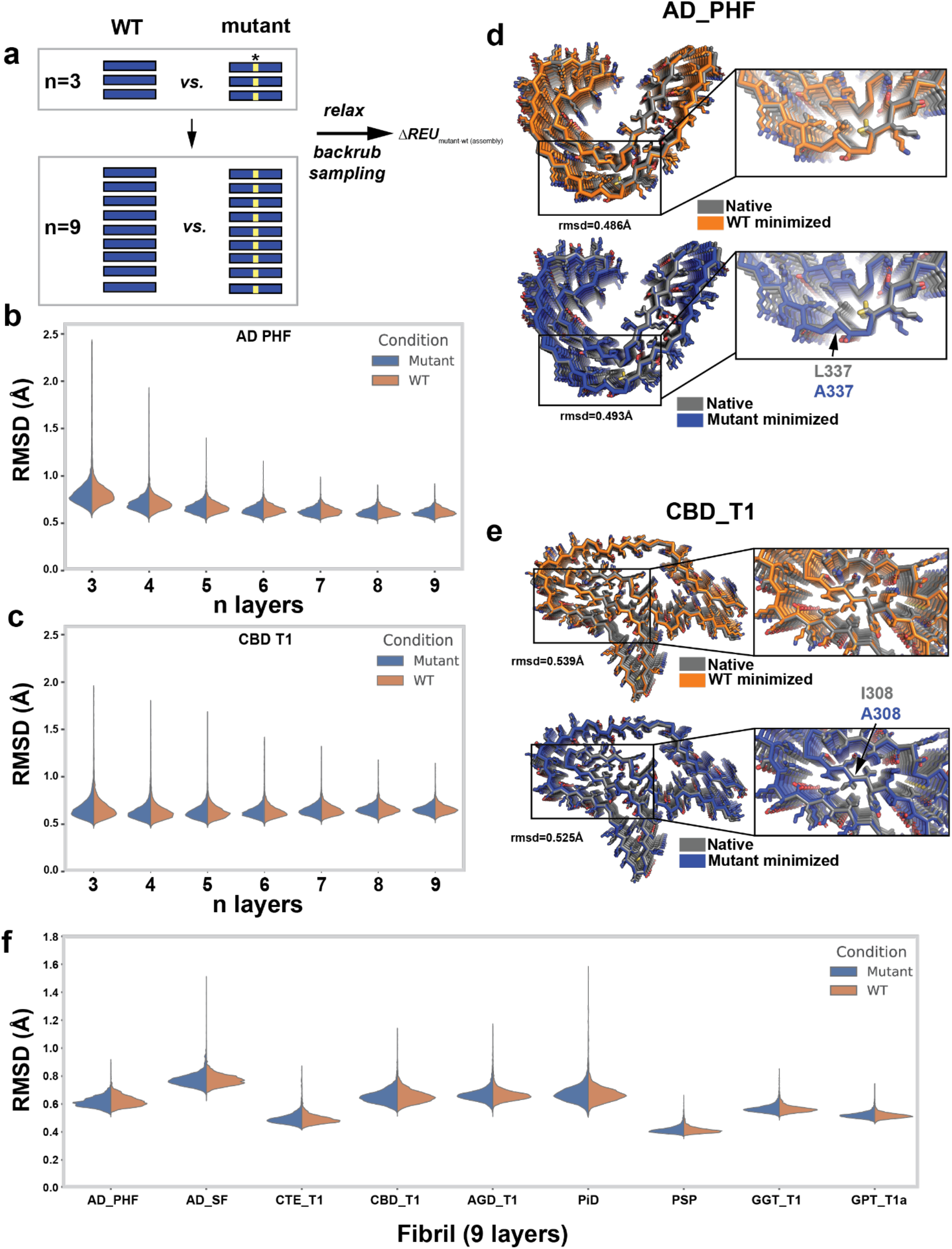
Fibril alanine scanning method produces minimized fibril assemblies that retain native contacts. **(a)** Schematic illustrating Flex ddG-based protocol to minimize and substitute alanine at individual positions in amyloid fibrils using different numbers of monomer layers. Blue box highlights a single layer. Yellow box highlighted by star denotes location of mutation. **(b-c)** RMSD distributions for WT (orange) and alanine mutants (blue) across different numbers of layers (n=3 to n=9) for AD-PHF (PDB id: 5o3l) **(b)** and CBD (PDB id: 6tjo) **(c). (d)** Structural overlay of AD-PHF nine-layer native WT (grey) and minimized WT (orange) reveals a RMSD of 0.486 Å (top). Structural overlay of AD-PHF nine-layer native WT (grey) and minimized L337A mutant (orange) reveals a RMSD of 0.493 Å (bottom). **(e)** Structural overlay of CBD nine-layer native WT (grey) and minimized WT (orange) reveals a RMSD of 0.539 Å (top). Structural overlay of CBD nine-layer native WT (grey) and minimized I308A mutant (orange) reveals a RMSD of 0.525 Å (bottom). **(f)** RMSD distributions for minimized WT (orange) and alanine mutant (blue) nine-layer structures reveals low RMSDs that range from 0.3 to 0.6 Å. The RMSD distributions are shown across 35 replicates at each position in each fibril and plotted as violin plots. AD-PHF, CBD_T1, CTE_T1, PiD, AGD_T1, PSP, GGT_T1a and GPT_T1 for PDB ids: 5o3l, 5o3t, 6gx5, 6nwp, 6tjo, 7p6d, 7p65, 7p66 and 7p6a.

Using the nine-layer fibril framework we minimized twenty-seven structures covering all known tau fibril structural polymorphs, including both patient-derived and recombinant fibrils, yielding structures that have near-native RMSDs (Fig. 3f). The starting fibril structures overall are not energetically favorable as determined by Rosetta but following iterations of minimization we produce low energy conformations for wild-type and all alanine mutant structures (Supplementary Fig. 3d). When comparing the wild-type and alanine mutant structures generated by this protocol, we find that overall, the alanine mutants are only slightly destabilizing, suggesting that alanine substitutions do not substantially alter the overall energy distributions when considering them cumulatively (Fig. 3f) or induce major deviation from the native conformation (Supplementary Fig. 3d). We find that the fibril structures with simpler topologies, namely the AD-PHF, AD-SF, and heparin-derived fibrils with only one-sided ^306^VQIVYK^311^ interactions, yield energies between −700 REU and −1200 REU while the more complex topologies (CBD, AGD, PSP, GPT and GGT) tend to yield lower energies around −1300 to −1600 REU (Supplementary Fig. 3d, bottom panel). Interestingly, the structures of recombinant fibrils induced with heparin yielded the highest energies (*i.e.,* least stable structures), likely because the interactions are not as well packed compared to the patient-derived conformations (Supplementary Fig. 3d, bottom panel). To explain this in more detail we normalized the energies to the number residues in the assembly (Supplementary Fig. 3d, top panel) and find that indeed the recombinant heparin-derived structures have some of the highest per residue energies. Interestingly, while the AD-PHF/AD-SF structures are similar to the CTE structures, we find that the CTE structures have equal or lower per residue contribution suggesting that the more open “C” conformation may be better packed than the more closed “C” from AD. Similarly, the other two-layer PiD structure has very low per residue energetics. Of the complex three-layer fibril topologies, PSP, GGT and GPT have the lowest per residue energies compared to AGD and CBD suggesting there is large variation in sidechain energetics of the more complex folds. Two of the CBD structures (PDB ids 6vh7 and 6vha) did not minimize as well, leading to higher per-residue energetics. This may be due to lower structure resolution causing slight conformational differences from the higher resolution, better minimizing CBD structures (PDB ids 6tjo/6tjx). We next evaluated in more detail whether alanine substitutions alter the fibrillar conformation and influence the energy of the structures. We find that in our backrub sampling protocol followed by alanine substitutions and minimization and does not significantly alter the overall RMSD distributions in the structures as compared to the native conformation (Fig. 3b, c). For example, the conformations of the minimized AD PHF L337A and the CBD I308A remain similar to the native structure with RMSD’s of 0.493 Å and 0.525 Å, respectively (Fig. 3d, e).

We found, however, that glycine to alanine substitutions often resulted in anomalously high structural energies. We note that glycine residues in all deposited (thirty-five total) tau fibril structures (red points, Supplementary Fig. 3e) adopt alternate torsional angles that are not often sampled in globular proteins (X-ray structures between 1.5 Å and 2.5 Å resolution). This observation is further extended to all deposited fibril structures (ninety-six) determined by cryo-EM using helical reconstruction in Relion (Supplementary Fig. 3f). We are unsure whether this is because the cryo-EM density maps are under determined and the energy of the model fitting is driven by empirical terms from force fields or whether these are *bona fide* features of the fibril structures. To see whether the backbone torsional angles for glycine residues are compatible with alanine substitutions we compared the glycine residue distributions in the tau fibril structures to observed alanine torsional angles in our globular protein dataset and find that they often are not permissible (Supplementary Fig. 3g).

As such, substitution of glycine residues in the fibril structures with alanine often yields large changes in energy dominated by the full atom Lennard-Jones repulsive potential energy component (herein fa_rep, representing Pauli repulsion of proximal atoms) suggesting the formation of van der Waals (vdW) atom clashes with a minor effect from the Dunbrack rotamer energy term (Supplementary Fig. 4a). Consistent with our original observations glycine to alanine substitutions contribute large increases to the fa_rep energy component likely through forming atom clashes but consequently also contribute a favorably to the full atom Lennard-Jones attractive potential energy term (herein fa_atr) (Supplementary Fig. 4a). Between this deviation in the backbone torsional angles and the energetic consequences of introducing atom clashes, we justify exclusion of glycine to alanine mutants from the analysis of energies. With the exception of these glycine to alanine mutants, however, our protocol appears to yield energetically favorable structures without large conformational deviations from the native.

As the nine-layer monomer stack preserves the native conformation in our protocol, we can investigate the native state interactions found in tau fibrils. Using mutagenesis as a probe, we proceeded to *in-silico* energetic analysis of twenty-seven patient-derived and recombinant structures of tau fibrils [15-18, 20, 21].

### *In silico* alanine-scanning captures the energetics of native fibril conformations

The Flex-ddG framework was originally implemented to capture the changes in binding energies for protein complexes. To adapt the Flex-ddG method to our fibrillar system we attempted two approaches. First, we probed the inter-layer energetics of the fibril by separating a central trimer from the rest of the nine-mer as the “unbound state” and compared its energetics to a “bound” state, where the central trimer occupies its native position in the fibril in order to probe the interface energy between layers of the fibril (Supplementary Fig. 4b). We find that the predicted ΔΔG*^interface^* (i.e., the ΔΔG*^binding^_mut-wt_* of the central trimer interacting with the remainder of the nine-layer fibril) has relatively small changes in response to alanine substitutions (Supplementary Fig. 4c, top panel). This suggests that the surfaces between the fibril monomers are relatively flat and do not have features that significantly stabilize the interactions beyond the backbone hydrogen bonding pattern which scales with the number of amino acids.

As the ΔΔG*^interface^* does not capture the energetic changes of mutation within layers of the fibril, only between layers, we next compared the difference in total energies of nine-mer assemblies between wild-type and alanine mutants *(i.e.*, the ΔREU*^assembly^_mut-wt_*), interpreting the total energy of the nine-mer assemblies and observe perturbations that are tenfold greater than the “bound” vs “unbound” interface energies (Supplementary Fig. 4c, bottom panel) and correlate with each other across the nine prototype structural polymorphs (Supplementary Fig. 4d). Although our approach cannot accurately estimate the energy of the unfolded ensemble of tau to predict a true ΔΔG*^mutation^*, this approach captures changes in both inter-layer and intra-layer energetics resulting from mutation, allowing us to interpret the contribution of an individual residue to the stability of the amyloid fibril.

We proceeded to use this second, total energy method for the analysis of twenty-seven tau structures using the nine-layer format, systematically substituting alanine residues at each position in the nine-layer stack and comparing the total energy to that of the wild-type assembly (Fig. 3a). From the “total energy” change between the wild-type and mutant assemblies, looking at the 9 distinct *ex vivo* tau protofilament structures we find that there is an alternating pattern consistent with β-sheets where residues facing inward contribute more than residues facing outward (i.e., solvent-facing residues) (Supplementary Fig. 4c, bottom panel). We parsed the different energy terms to understand the origins of the energy differences. On inspection of the energy changes for different amino acid types upon mutation to alanine, we find that non-polar amino acids (Ile, Val, Leu, Phe, Tyr and Met) have the largest loss in the fa_atr energy term and some gain in the fa_rep energy term, consistent with creating destabilizing cavities (Supplementary Fig. 4a) in the cores of fibrils (Supplementary Figs. 2 and 3a). We also observe that the fa_atr energy term for nonpolar residues contribute similarly across each of nine *ex vivo* fibril structures (Supplementary Fig. 5a). Lastly, we interpret the solvation energy term and find that on average alanine substitution of non-glycine and nonpolar residues yield favorable solvation energetics for all nine structures in aggregate (Supplementary Fig. 4a) and also individually (Supplementary Fig. 5b). This term indeed offsets some of the destabilizing contribution of mutating nonpolar residues to alanine but is relatively minor suggesting that formation of the void and loss of vdW contacts dominates the energy change of nonpolar to alanine mutants. These data for the first time uncover the general folding and energetic principles of tau assemblies and relate these principles to the possible first steps in monomer folding, which may be defined by burial of nonpolar amino acids and arrangement of lysine residues on the fibril surface.

### Identification of thermodynamic hotspots of aggregation

We next interpret energetics to understand on a per residue and motif level to gain insight into the rules that govern stability in fibril structures. To more easily visualize regions that may be important for stabilizing fibrils we calculated an average change in energy (ΔREU*^assembly^_mut-wt_*) per residue for a five-residue window. We relate these values to amyloid propensity and find that many of the interactions between energetically important elements involve amyloidogenic segments including ^275^VQIINK^280^, ^306^VQIVYK^311^, ^337^VEVKSE^342^ and others (Fig. 4a). Indeed, these interaction hotspots also correlate to fragments that encode clusters of nonpolar residues (Fig. 4b). We also map the per residue energy change of mutation (ΔREU*^assembly^_mut-wt_*) onto the nine distinct *ex vivo* fibril protofilaments and unsurprisingly observe that residues that contribute to the largest change in energy are often buried in the fibril core and that residues on the periphery tend to have smaller energetic contributions (Fig. 4c). Consistent with this observation we find alanine mutations significantly destabilize the fa_atr term, indicating that burial of nonpolar residues in tau is a dominant contributor to fibril stability (Supplementary Fig. 4a). Additionally, we find that the Lazaridis-Karplus solvation energy component (fa_sol) term generally decreases (i.e., becomes more favorable) with alanine substitutions because this allows favorable interactions with solvent, but these gains in solvation energy are often balanced with loss in vdW contacts. We fit the relationship between per residue change in stability to ΔSASA*^folding^ _single layer_* and ΔSASA*^folding^ _fibril_* as a function of amino acid types. We see that as expected, all amino acid types experience some degree of change in SASA upon folding into monomer conformation (*i.e.*, as a single layer of the fibril), and then a further second stage of burial when that folded monomer conformation is incorporated as a layer of a fibril. Overall, the two terms cluster residues together, but the coefficients of determination (R^2^) are low, suggesting that each residue type has a significant number of outliers that cannot be explained by the fit. For all residue types, there are some residues with high ΔSASA*^folding^ _single layer_* and ΔSASA*^folding^ _fibril_*, but low impact on fibril stability as measured by ΔREU when mutated to alanine. This may be due to inwards facing residues in the fibril that face voids, such as those in the CBD structures. We find that for both ΔSASA*^folding^ _single layer_* and ΔSASA*^folding^ _fibril_*, the nonpolar residues have the largest baseline contribution in change in energy per change in SASA, indicating that their burial is a major factor associated with fibril stability (Supplementary Fig. 5c). Both acidic and basic residues have populations with low change in energy when mutated to alanine but there is a subpopulation with both a large ΔSASA*^folding^* and large change in energy. This indicates that there exists a set of buried charged residues which are important contributors to fibril stability. Indeed, there are several buried salt-bridges that involve lysines and acidic residues that account for this behavior. For some fibril structures, including Huntingtin or CPEB protein, a large portion of buried contacts involve polar amino acids [52, 53]. In our tau fibril analysis, it appears that nonpolar residues are largely buried in the structures leaving mostly polar and specifically basic residues on the surface with some exceptions (Supplementary Fig. 2c and Supplementary Fig. 3a). Indeed, the unique protofilaments which have been identified across diseases leverage these surface residues to adopt alternate protofilament arrangements stabilized by the weak polar interactions that extend along the fibril axis (e.g., the Alzheimer’s Disease PHF/SF or the GGT type 1/2/3 fibrils).

**Figure 4.**
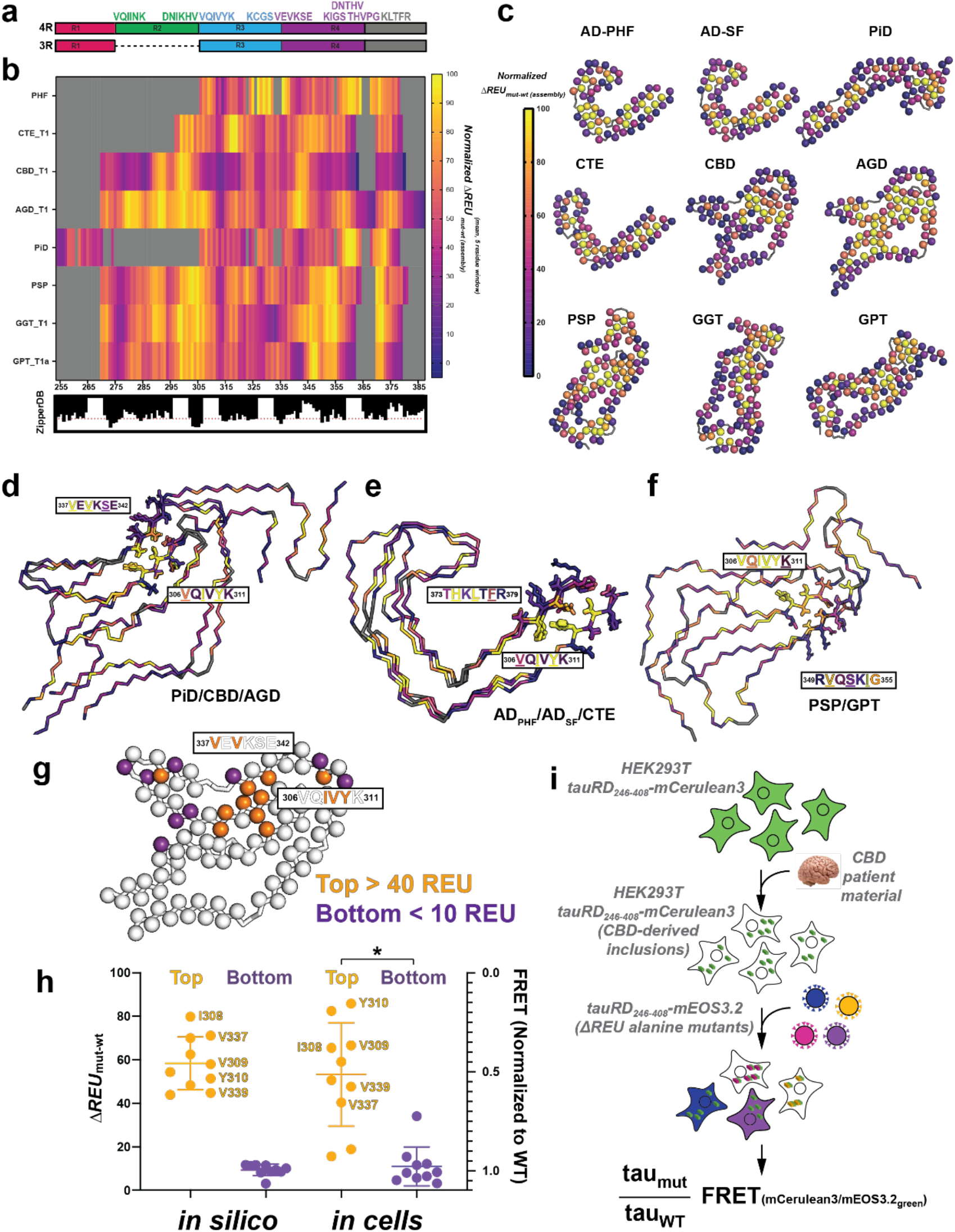
Fibril alanine scanning uncovers modular networks of hotspot interactions in tau fibrils. **(a)** Cartoon illustration of 4R and 3R isoforms of the tau repeat domains colored as in Fig. 1A highlighting location of key amyloidogenic motifs shown above. **(b)** Normalized heatmap of energetic change in response to substitution to alanine across a 5-residue window for 9 prototypical tau fibril structures: AD-PHF, CBD_T1, CTE_T1, PiD, AGD_T1, PSP, GGT_T1a and GPT_T1. PDB ids: 5o3l, 6gx5, 6nwp, 6tjo, 7p6d, 7p65, 7p66 and 7p6a. Scale is colored in plasma from yellow (important) to purple (not important). **(c)** Mapping per residue ΔREU^assembly^_mut-wt_ profiles onto the 9 tau fibril structures uncovers inward facing amino acids contribute more to stability than surface exposed residues. Structures are shown as a single layer shown in ribbons with C-α atoms shown in spheres for each amino acid. The C-α is colored by changes in energy due to individual amino acid mutations to alanine using the plasma color scheme, yellow (100 normalized ΔREUs, important) to purple (0 normalized ΔREUs, not important). **(d-f)** Structural overall of similar structures highlighting energetically important interactions with VQIVYK. **(d)** Overlay of PiD, CBD and AGD highlighting the energetically important and conserved interactions between ^306^VQIVYK^311^ and ^337^VEVKSE^342^. (**e**) Overlay of AD-PHF, AD-SF and CTE highlighting the energetically important and conserved interactions between ^306^VQIVYK^311^ and ^373^THKLTFR^379^. **(f)** Overlay of PSP and GPT highlighting the energetically important and conserved interactions between ^306^VQIVYK^311^ and ^337^VEVKSE^342^. The fibril structures are shown as a single layer in ribbon representation, the interacting motifs are shown as sticks and colored by their normalized per residue energy contribution using the plasma color scheme (yellow=100 and purple=0). (**g**) Cartoon schematic to evaluate the importance of key amino acids in cellular tau seeding experiments using CBD patient tissues. Orange residues are predicted to be important to fibril stability (normalized ΔREU^assembly^_mut-wt_ > 40), while purple residues are predicted to be minimally important to fibril stability (normalized ΔREU^assembly^_mut-wt_ < 10). (**h**) Plot comparing the top-10 and bottom- 10 most important residues as determined by ΔREU^assembly^_mut-wt_ from the *in silico* alanine scan (scaled from normalized 0 ΔREU^assembly^_mut-wt_, *i.e.*, no predicted loss in fibril stability caused by alanine mutant to normalized 100 ΔREU^assembly^_mut-wt_ - high predicted loss in fibril stability due to alanine mutant), alongside the effect of these alanine mutants on the ability of Corticobasal Degeneration (CBD)-brain derived tau seeds to seed aggregation as measured by the biosensor-based aggregation assay. (Range from 1.0=no change in FRET, *i.e.*, no effect on ability of mutant tau to aggregate in biosensor cells when seeded with CBD-derived seeds as compared to the WT, to 0.0=no FRET observed, *i.e.*, mutant tau is unable to induce FRET-detectable aggregation in the biosensor). The data is shown as averages with standard deviation. The experimental top and bottom hits are shown to be significantly different with p-values <0.0001 using a two-tailed Mann-Whitney test and are indicated by a star. (**i**) Cartoon schematic showing the method of the *in vitro*, FRET biosensor-based tau seeding cell assay used to assess impact of mutants on tau’s ability to aggregate when seeded by proteopathic tau seeds.

To understand the energetics of the ^306^VQIVYK^311^ motif specifically, we looked at the energetics of the residues identified to interact with the amyloid motif in the different structures (Fig. 1). We find that in the PiD, CBD and AGD structures, V306, I308 and Y310 from ^306^VQIVYK^311^ make stabilizing interactions with V337 and V339 from ^337^VEVKSE^342^ (Fig. 4d). Similarly, we find that in the AD-PHF, AD-SF and CTE structures, V306, I308 and Y311 of the ^306^VQIVYK^311^ contribute stabilizing interactions with L376 and F378 of ^373^THKLTFR^379^ (Fig. 4e). In the PSP and GPT structures, Q307, V309, K311 of VQIVYK form stabilizing interactions with V350, S352 and I354 of ^349^RVQSKIGS^355^ (Fig. 4f). Additionally, we find that in CBD, AGD (and GGT) there are hotspot interactions between ^306^VQIVYK^311^ and ^295^DNIKHV^300^ mediated by nonpolar contacts centered on V306 and I297 (Supplementary Fig. 6a). Excitingly, this element was consistently formed in a core of fibril structures assembled from a recombinant fragment and is similar to the core element in the GGT structure [49]. These data suggest that ^306^VQIVYK^311^ plays important, stabilizing roles in diverse structural polymorphs observed in different diseases. The VQIVYK-based networks of interactions may help determine the fold and govern whether the fibril adopts local interactions with the ^295^DNIKHV^300^ sequence in two different ways (CBD, AGD vs PSP, GPT), but also help establish long range contacts to ^337^VEVKSE^342^ (CBD, AGD and PiD) or ^373^THKLTF^378^ (AD-PHF, AD-SF and CTE). While ^306^VQIVYK^311^ seems to play a central role in all *ex vivo* (and nearly all recombinant with the exception of RNA-induced tau fibrils) fibrils, we also observe features that utilize other elements such as conserved and energetically stabilizing interactions between ^355^DNITHV^363^ and ^370^KKIETH^376^ in the GGT, PSP and PiD structures (Supplementary Fig. 6b) [54]. These interactions are peripheral to the central cores but are related to the ^295^DNIKHV^300^ and ^306^VQIVYK^311^ interaction because they similarly surround a β-turn capable PGGG motif. Understanding how these contacts are formed may help us facilitate controlling the order of interactions that lead to a fold observed in disease.

To test whether the aggregation-prone ^306^VQIVYK^311^ peptide alone can engage in heterotypic interactions with other nonpolar elements predicted from the hotspot calculations, we designed co-aggregation experiments using ^306^VQIVYK^311^ with ^337^VEVKSE^342^ and ^350^VQSKIG^355^, two different VQIVYK-interacting elements predicted to stabilize the different fibril structures. We used a ThT fluorescence aggregation assay and then compared endpoint fluorescence signal to understand whether any of these inert peptides can regulate assembly of ^306^VQIVYK^311^. We performed aggregation of the ^306^VQIVYK^311^ peptide at 200 µM alone, and in the presence of each heterotypic contact peptide at 200, 100, and 50 µM. We also measured the ThT signal of the test peptides at the three concentrations and confirmed the presence or absence of fibrils in all samples by TEM. (Supplementary Fig. 6c-f). We find that each peptide influences ^306^VQIVYK^311^ aggregation in different ways. The most dramatic effect was observed with ^350^VQSKIG^355^ (Fig. 4f; PSP and GPT) which blocked ^306^VQIVYK^311^ aggregation completely (Supplementary Figs. 6c). The ^337^VEVKSE^342^ peptide (Fig. 4e; CBD, PiD and AGD) only inhibited VQIVYK aggregation at the highest concentration (200 µM) (Supplementary Figs. 6d). To further probe interactions between VQIVYK and VEVKSE, we also coaggregated VQIVYK with VEVKSE and two mutants of VEVKSE at hotspot positions mutating each valine to alanine. We also included an experiment with a soluble control peptide (sequence GSPGGS) with a proline intended to reduce β-sheet propensity. We find that VEVKSE yielded a similar decrease to that seen in the previous experiment, while the control peptide had no effect (Supplementary Fig. 6e). Interestingly, the AEVKSE peptide was not able to efficiently inhibit VQIVYK aggregation while VEAKSE was even more efficient (Supplementary Fig. 6e). These data suggest that leveraging peptide aggregation systems it might be possible to regulate amyloid motif aggregation.

Leveraging the capacity of tau to propagate seed conformations in cellular systems, we wanted to test residues that we predict are energetically important for the stability of the CBD tau fibril structure in cells. Expression of WT tau as a fusion to a FRET compatible fluorescent reporter (*e.g.*, mClover) can be used to induce aggregation of the intracellular reporter by transfection of a recombinant or a tauopathy patient-derived seed [55]. Subsequently, tau fused to a FRET compatible pair (mCerluean, Ruby, etc.) can be transfected and the incorporation of the new reporter into the existing aggregates measured by FRET. More recently, this system has been shown to replicate the conformation of tau seeds from cells into animal models and back into cells [56]. We used this system to test our top ten ΔREU^assembly^_mut-wt_ predicted hits (Fig. 4g,h, orange) identified in the CBD *in silico* alanine scan and as a control we also tested ten neutral ΔREU^assembly^_mut-wt_ hits (Fig. 4g,h, purple). Consistent with our observations, the top hits from our calculations cluster to the contacts between the ^306^VQIVYK^311^ and ^337^VEVKSE^342^ motifs (Fig. 4g). We produced cell lines expressing WT tau fused to mClover3 which were treated with CBD patient material to replicate the CBD conformation in the cells. Subsequently, we transduce tau fused to mEOS3.2 [57] encoding alanine mutations to test the ability of each mutation to incorporate into the WT propagated CBD-derived inclusions in the cells. Mutations that interfere with incorporation into CBD-derived inclusions should not yield FRET between mCerulean3 and mEOS3.2 while mutations that can incorporate should have similar FRET to WT tau. To estimate the effect of these mutations, we compare the FRET signal of the mutant to the WT tau (Fig. 4i and Supplementary Fig. 4f). We find that the experimental data compare well with the ΔREU^assembly^_mut-wt_ calculations and similarly allow separation of the top hits from the bottom hits with significant p-values (Fig. 4h). For the top hits we predict correctly 8 of 10 positions which cluster to the important amyloid motif interactions (Fig. 4h). The 2 predicted positions at F346A and P312A that do not match the experiment, fall outside of the core of interactions, highlighting potential failure of the energy function to capture the energetics of these contacts correctly or inability of our method to model larger conformational shifts that may be caused by mutation. Our experiments leverage tau’s capacity to replicate fibril conformations in cells and allow us to experimentally validate our computational method to discover hotspot residues important for fibril stability.

Through computational alanine scanning, we can identify hotspot residues and sequence regions that form the heterotypic interactions that stabilize fibril structures. We observe these interactions in *in* vitro aggregation experiments and validate the importance of hotspot residues to fibril formation using an in-cell model of tau amyloid aggregation. We anticipate that stabilizing these interactions within a unique tau fold while destabilizing them in other structures will restrict the tau monomer to adopt only that single tau fibril fold.

### Classification of structures and their features based on their energetic profiles

We next employed a machine learning approach to begin classifying the structures based on the per residue ΔREU^assembly^_mut-wt_ values to learn what residues/features may help cluster and discriminate fibril conformations. As some of the structures differ in sequence composition, we first curated the ΔREU^assembly^_mut-wt_ data to only contain residues present in all of the structures. For example, PiD fibrils are comprised of only a 3R tau isoform lacking the second repeat domain, AD/CTE are limited to repeat domains three and four, while others (AGD, CBD, GGT, PSP, and GPT) contain sequence from repeat domain two, three, and four. From this reduced set of common residues, we calculated a distance matrix of all fourteen disease-derived tau fibril polymorphs, which include subtypes for some of the structures including AD, AGD, GGT and GPT (Supplementary Fig. 7a). This distance matrix was then used to construct a hierarchical clustering of the different fibril types (Fig. 5a and Supplementary Fig. 7b). We find that related subtypes cluster together and reassuringly structures such as AD-PHF, AD-SF and CTE or AGD_T1, AGD_T2 and CBD cluster together. Likewise, the GPT and GGT structures cluster together with PSP. Again, this observation is consistent with the GPT structure being described as a hybrid PSP and GGT structure. Consistent with our prior observations these more related structures also contain conserved interactions involving ^306^VQIVYK^311^. Interestingly, the PiD structure clusters together with GPT. This anomalous clustering of PiD is perhaps not surprising because of this conformation’s unique sequence composition, involving residues from the first repeat domain. As such, a portion of the sequence is unique and cannot be used in clustering and structure classification with this method. After identification of common features, we focused our efforts to tease out more subtle differences between related structures which would be difficult to discriminate via the surface using proteins (i.e., antibodies). Being able to predict residues that form energetically discerning interactions may allow the design of tau sequences at specific sites to stabilize one conformation while destabilizing another, allowing the creation of designer tau sequences that can only propagate a single conformation while being incompatible with others.

**Figure 5.**
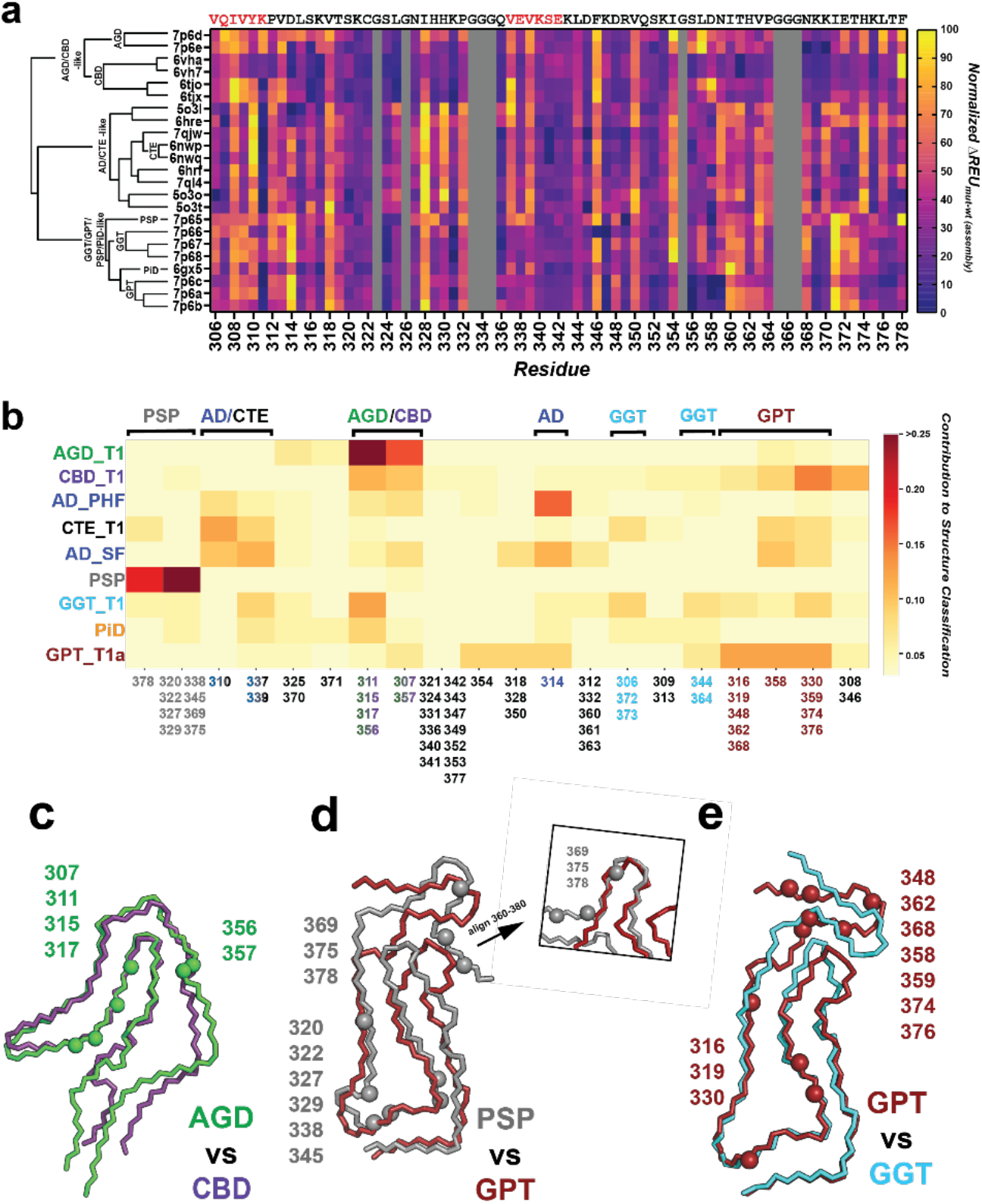
Feature classification clusters structural polymorphs by hotspot interactions and folds. **(a)** Dendrogram produced by hierarchical/agglomerative clustering via Ward’s method of tau fibril structures based on normalized *in silico -* ΔREU^assembly^_mut-wt_ values determined by the alanine scan, displayed next to plot of normalized ΔREU^assembly^_mut-wt_ values. Scale is colored in plasma from yellow (highest *in silico* ΔREU^assembly^_mut-wt_) to purple (lowest *in silico* ΔREU^assembly^_mut-wt_) for all available tau *ex vivo* fibril structures from AD, CBD, AGD, PSP, GPT, GGT and PiD, including subtypes. PDB id: 7p6d, 7p6e, 6vha, 6vh7, 6tjo, 6tjx, 5o3l, 5o3t, 6hre, 7qjw, 6nwp, 6nwq, 6hrf, 7ql4, 5o3o, 5o3t, 7p65, 7p66, 7p67, 7p68, 6gx5, 7p6c, 7p6a and 7p6b. **(b)** Contribution of residues/groups of residues towards correct classification of an aggregate structure by a random forest classification model trained on the *in silico* ΔREU^assembly^_mut-wt_ of alanine mutants. Clusters most important for correct identification of a structure are colored red, while the least important clusters for a structure’s identification are colored pale yellow. Residue labels are colored by the structure type they most contribute to identification of-CBD (purple), AGD (green), AD PHF/SF (blue), CTE (black), PSP (grey), GGT (cyan), PiD (orange), or GPT (red). Less important residue clusters for classification/identification are colored light grey. **(c-e)** Overlaid fibril monomers of similar structure with residues that most contribute to their classification shown with C-α atom spheres. **(c)** Overlay of CBD (blue) with AGD (green), highlighting CBD residue 358 with C-α atom shown as a blue sphere and AGD residues 311, 315, 317, 356 C-α atoms shown as green spheres. **(d)** Overlay of PSP (grey) with GPT (red), highlighting PSP residues 307, 320, 322, 327, 329, 338, 351, 375, 378 with C-α atoms shown as grey spheres and GPT residues 319, 348, 354, 359, 362, and 376 C-α atoms shown as red spheres. **(e)** Overlay GPT (red) with GGT (cyan), highlighting GPT residues 319, 348, 354, 359, 362, and 376 C-α atoms shown as red spheres and GGT residues 354 and 357 C-α atoms shown as cyan spheres.

### Identification of differentiating features among disease-associated tau fibrils

We proceeded to identification of distinguishing features between tau fibrils by training and interpreting a machine learning model on the per-residue ΔREU^assembly^_mut-wt_ values generated by the *in silico* fibril alanine scan. Briefly, we performed hierarchical clustering on the residue ΔREU^assembly^_mut-wt_ values and noted that many sets of residues had highly similar behavior between fibrils (Supplementary Fig. 7b). To reduce data dimensionality and simplify interpretation of the classification model by leveraging this covariance of residues, we proceeded to perform feature agglomeration, recursively combining covarying groups of residues into single features (Supplementary Fig. 7c). With this reduced set of features, we trained a random forest classification model to classify the various disease associated fibrils and subtypes into their parent fibril types (e.g., GGT_T1/T2/T3 as GGT, AD-PHF/SF as AD, GPT_T1a/T1b/T2 as GPT, etc.). 2500 classification models were generated using 25% of the replicates generated by the alanine scanning protocol as training data and the remaining 75% of the replicates as testing data to verify performance of the classification model, selecting the model with the highest classification accuracy on the testing set. A model was selected with >99.5% prediction accuracy on the test set, showing ability to effectively discriminate structures based solely on their ΔREU^assembly^_mut-wt_ values. Despite its highly accurate classification performance, the model is least confident in its classification of AD-PHF/AD-SF vs CTE_T1 fibrils (Supplementary Fig.7d). This is likely due to the high degree of mutual similarity of the AD-PHF, AD-SF, and CTE_T1 structures (and thus the ΔREU^assembly^_mut-wt_ values derived from them).

Subsequently, this model was used to identify major distinguishing features of each disease fibril type by running the classifier model on the per-residue ΔREU^assembly^_mut-wt_ values (mean of all thirty-five replicates) and inspecting the model to see to which degree each residue cluster contributed to the correct classification of each structure (all structures were correctly classified by the model) (Fig. 5b). Notably, this reveals clusters/sets of residues that have unique behavior in certain fibrils/sets of fibrils as discerned by the model and used in identification of the structure. We find that residues Q307, K311, L315, K317, S356, and L357 are strongly distinctive in AGD/CBD. We see weaker contribution of single clusters of residues to prediction of AD vs CTE fibrils, again suggesting that differences between the two are relatively minor and forcing the model to parse out smaller differences over a larger number of fibrillar features to make an accurate classification. We map these values onto the structures and note that these discriminative positions highlight locations where fibril structures deviate from each other. For example, AGD-discriminating residues lie where the residues ^312^PVDLSK^317^ interact ^285^SNVQSKGS^293^, a region with differing backbone geometry between AGD and CBD. Similarly, residues S356, and L357 are also located in a region that deviates between AGD and CBD (Fig 5c). Taking the AD/CTE fibrils together, differentiating residues here highlight the residues differing in the one-vs-two-sided ^306^VQIVYK^311^ interaction that separates AD, CTE and PiD from the other fibrils. These discriminative residues also localize to regions of diverging structure between PSP/GPT and GGT/GPT, suggesting the classification model is successfully identifying regions of distinction between tau fibril structures solely based on the per-residue ΔREU^assembly^_mut-wt_ values. We anticipate that accurate identification of distinguishing regions of tau fibrils can contribute to the design and development of tools to distinguish fibril conformations (e.g., biosensors and antibodies) and engineered protein sequences that more readily adopt desired fibril conformations, as well as helping lend insight into the differing disease processes that produce these unique conformers.

## Discussion

We have developed an *in silico* alanine scanning method to interpret the energetic contribution of amino acids in fibrillar structures obtained using cryo-EM methods. Our work presents the first comprehensive analysis of cryo-EM tau fibril structures that uncovers amino acids (i.e., hotspot residues) that stabilize intra-peptide interactions in the distinct structural polymorphs. These hotspot residues are often nonpolar and lie in sequences that are either amyloidogenic or interface with amyloidogenic motifs. Interestingly, we discover that these residues form interaction motifs that are often modular, being combined together in various ways across the different fibril structures (Figure 6). Related fibril conformations use similar sets of these hotspot interactions. This modularity of interactions uncovers the underlying interactions that enable the different fibrillar folds in various structural polymorphs. We use cellular and *in vitro* aggregation experiments to validate energetically important interactions predicted by our methods. We also used our calculations to classify diverse tau fibril structures and uncover sets of modular interaction motifs that are key features which permit clustering of related structures and conversely differentiate distinct folds. These uncovered interactions mirror our initial observation surrounding the ^306^VQIVYK^311^ motif. We anticipate that our method will be broadly useful for the interpretation of folding and interaction energies in fibrils structures and acknowledge that future work must focus on developing experimental methods to capture these energetics-either directly or by proxy- to improve our predictions. These data will be important to interpret how pathogenic mutations alter the conformation of proteins and set the stage to begin designing proteins sequences that preferentially adopt unique structural folds.

**Figure 6.**
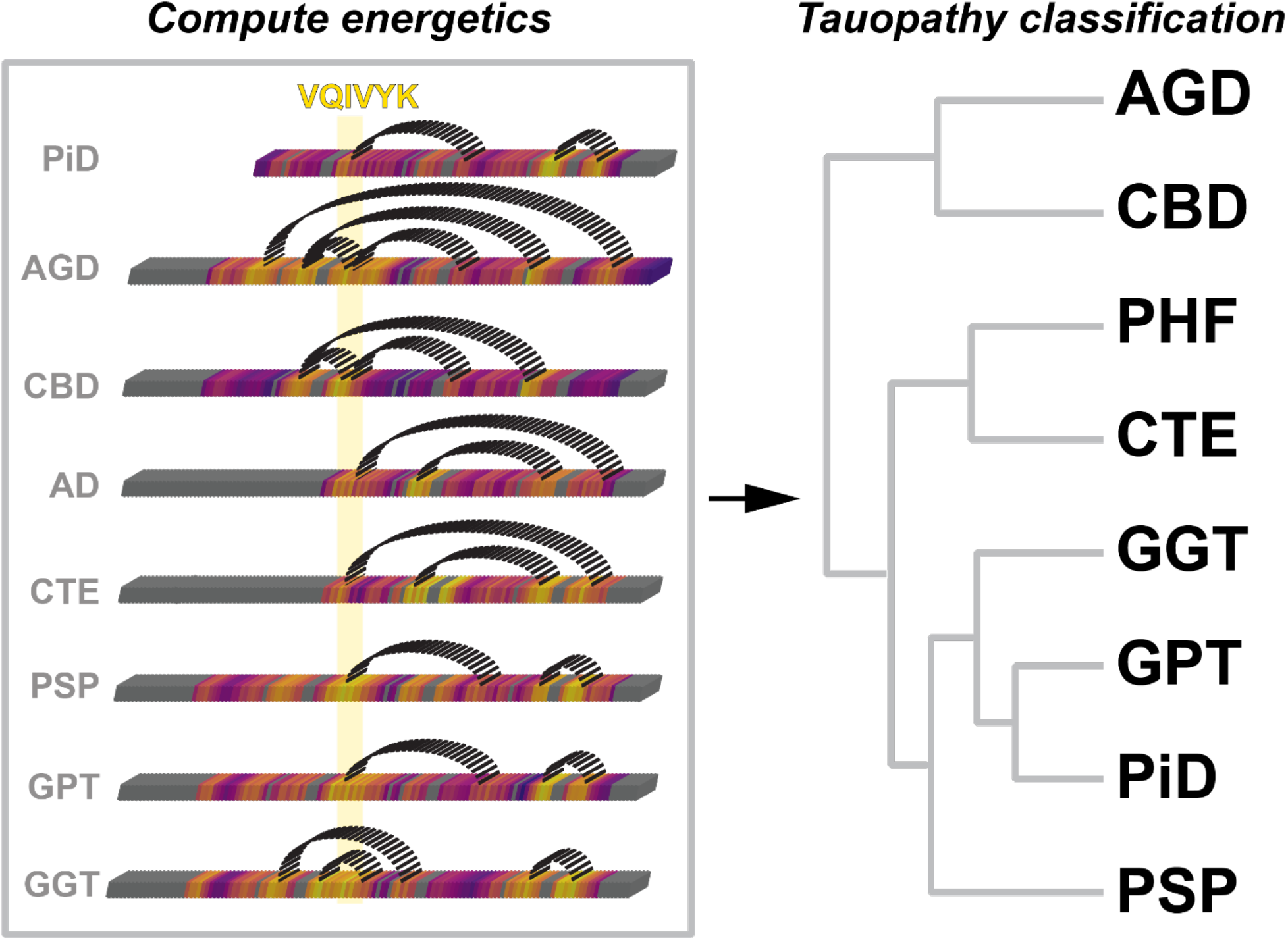
Classification of tau fibril structures using per residue energetics. Contacts observed in fibrils between energetically important hetero-typic interactions determine tauopathy folds. The linear heatmaps are shown as 3D rendered boxes that are colored by the normalized change in energy values (yellow=stabilizing and purple=neutral). Dashed semi-circles link contacts between hotspot regions that interact in the different structures are used to classify the different folds.

### Diverse topologies assembled from simple rules

We present the first in-depth analysis of the large set of diverse tauopathy fibril structures. The evidence is emerging that distinct tau fibril structures are associated with different diseases. To date it remains unknown how these disease structures form; recombinant reproduction of disease fibrils remains a challenge, highlighted by the stark differences between *ex vivo* patient conformations compared to the heparin-derived recombinant forms. While understanding how these structures are formed remains at the forefront of identifying disease mechanisms and the development of conformation specific reagents to diagnose and treat tauopathies our analysis uncovers exciting rules that underlie the modularity of contacts in the fibril structures. We find the amyloid motifs in the structures play essential roles in promoting aggregation but also stabilizing contacts that determine each topology. Furthermore, the ^306^VQIVYK^311^ sequence has an intrinsic capacity to interact with itself and with other sequence encoding nonpolar residues. It remains unknown why this sequence is such an efficient aggregation promoter, but it likely is defined by the amino acid composition as well as its ability to adopt β-sheet conformations in solution allowing other nonpolar residues to engage in contacts with specified geometries on two surfaces. We see that removal of nonpolar residues important in these contacts impacts amyloid formation via both *in vitro* peptide aggregation and in cell fibril incorporation. An alternative approach to understanding how to produce recombinant tau structures that mimic disease fibrils is to reverse engineer the sequences to promote formation of substructures that recapitulate those in *ex vivo* fibrils. Recent successful efforts to fibrillize a recombinant fragment of R3R4 tau into AD and CTE conformations limit the possible interactions via use of a sequence fragment. Inclusion of specific anions/cations in the reaction mixture may allow further reduction of possible conformations by stabilizing nonpolar contacts. However, more complex topologies with longer sequences (and thus more possibilities for formation of different polymorphs) will likely remain a challenge to recapitulate in vitro [49]. It is possible in physiologic systems that other factors such as ligands or post-translational modifications such as phosphorylation, ubiquitination or acetylation may stabilize certain intermediate conformations to yield the formation of distinct fibril folds. Understanding the network of interactions responsible for the stability of fibrils may also provide insight into the physiological states and structural intermediates necessary to stabilize certain interactions and thus preferentially form particular fibril polymorphs.

### Reverse engineering structural elements to build fibrillar folds

We compare the current explosion of cryo-EM fibril structures to the early efforts of X-ray crystallography to determine structures of globular proteins. These initial efforts to determine structures of globular proteins created the structural knowledge that can now be used to develop improved algorithms to accurately predict protein structure from sequence and to design de novo protein structures with novel functions. We anticipate that as the cryo-EM fields focus on structure determination of fibrils matures, new computational methods will be developed that will accurately predict how sequences can convert from disordered ensembles of conformations to discrete fibrillar folds or ensembles of possible fibrillar conformations. Further advances in structure determination and protein modelling will begin to explore how one sequence can adopt an ensemble of different conformations and the requirements to understand rate limiting interactions that determine the formation of structural folds. We envision that these methods will help predict how sequence changes in a protein that has the capacity to adopt alternative fibrillar conformations may restrict the formation of distinct structures. There may be utility in designing more restrictive sequences that allow detection and differentiation of these structures, using cellular systems similar to biosensors developed for tau and other proteins that form amyloids and can replicate in a prion-like fashion. These modified sequences that can only adopt a single conformation may be used to detect a tauopathy-specific conformation thus leading the way to develop diagnostics to detect formation of disease-specific seeds prior to the accumulation of pathology.

## Conclusions

Together our experiments and computational method have uncovered how energetically important interactions that recur across different structural polymorphs may contribute to folding of tau into distinct structural polymorphs. These methods will help uncover the folding mechanisms of tau and other proteins into conformations associated with disease. Our energy calculations can be used to pinpoint energetically distinguishing interactions thus allowing the design of sequences that can propagate only one fibril conformation that may be useful for development of diagnostic reagents. Finally, our computational method is generalizable to any fibrillar structure and could be used to uncover energetically important contacts to begin classifying structures by their stabilizing interactions, as well as identifying more general patterns in amyloid aggregation and fibril structure within the entire amyloid proteome.

## Materials and Methods

### Amyloid motif prediction in tau and *in vitro* ThT peptide fibrillization

We used the ZipperDB database to identify possibly amyloidogenic motifs in tau [46]. Hexapeptides that showed REU below −25 were selected for subsequent in vitro aggregation experiments. C-terminal amidated and N-terminal acetylated hexapeptides representing regions of amyloidogenic behavior (as predicted by ZipperDB) were sourced from GenScript at >= 95% purity. The peptides were disaggregated in 300 μL of trifluoroacetic acid (TFA) and incubated at 30°C with 850 RPM orbital shaking for one hour. After disaggregation, peptides were blown dry of TFA under N_2_ gas and then lyophilized for two hours to remove residual TFA. For fibrillization, lyophilized peptides were resuspended in 500 μL of ultrapure water, vortexed thoroughly to resuspend, and then diluted (based on initial mass) to a final concentration of 1 mM peptide in 1X phosphate-buffered saline (PBS) using 50 μL of 10x PBS + NaOH (1.37 M NaCl, 27 mM KCl, 100 mM Na_2_HPO_4_, 18 mM KH_2_PO_4_, 1.56 μM NaOH) and ultrapure water. pH was verified to be neutral using pH test strips. Fibrilization was carried out for one week at 37°C with 900 RPM orbital shaking. Samples were taken from the in vitro fibrilization reactions for ThT measurement in 6-plicate. In the dark, 3 μL of 250 μM Thioflavin T (ThT) solution was mixed with 27 μL of the in vitro fibrilization reaction mix for a final ThT concentration of 15 μM. Six blank samples containing only PBS with 15 μM ThT were also prepared. The samples were loaded onto a clear-bottom, 384 well plate and read in a Tecan Infinite M100 at 446 nm excitation wavelength (5 nm bandwidth), 482 nm emission wavelength (5 nm bandwidth). The instrument was heated to 37°C, and samples were shaken for 10 seconds prior to acquisition of the data. The mean blank sample fluorescence was subtracted from the fibrillized peptide fluorescence and reported as the ThT signals for the samples.

### *In vitro* ThT aggregation of VQIVYK alanine mutant and co-aggregation of VQIVYK with competing peptides

For heterotypic VQIVYK co-aggregation experiments and aggregation of VQIVYK alanine mutants, all peptides were monomerized in TFA (as above). Reactions containing sufficient peptide for 350 μL of 200 μM VQIVYK alone, 200 μM VQIVYK alanine mutants, 200 μM VQIVYK with 50 μM, 100 μM or 200 μM VEVKSE, VQSKIG, AEVKSE, VEAKSE, GSPSGS peptides, or 200 μM VEVKSE, VQSKIG AEVKSE, VEAKSE or GSPSGS peptides alone were mixed and then blown dry of TFA under N_2_ gas and then lyophilized for two hours to remove residual TFA. In the dark, a 1X PBS + 25 μM ThT solution was prepared. Reactions were prepared by resuspending the mixed, lyophilized peptides in 350 μL 1X PBS + 25 μM ThT. The samples were loaded in six replicate, 50 μL reactions along with a 6, equally sized blank reactions containing only 1X PBS + 25 μM onto a clear-bottom, 384 well plate and read in a Tecan Infinite M100 at 446 nm excitation wavelength (5 nm bandwidth), 482 nm emission wavelength (5 nm bandwidth). The instrument was heated to 37°C, and samples were shaken for 10 seconds prior to acquisition of each data point. Data points were collected every 5 minutes for the first 8 hours and then every 30 minutes afterwards for one week and the blank fluorescence subtracted.

### Transmission Electron Microscopy

An aliquot of 5 μL sample was loaded onto a glow-discharged Formvar-coated 200-mesh copper grids for 30 s and was blotted by filter paper followed by washing the grid with 5 μL ddH_2_O. After another 30 seconds, 2% uranyl acetate was loaded on the grids and blotted again. The grid was dried for 1 minute and loaded into a FEI Tecnai G2 Spirit Biotwin TEM. All images were captured using a Gatan 2Kx2K multiport readout post column CCD at the UT Southwestern EM Core Facility.

### *In Silico* Rosetta ΔΔG^interface^ and ΔREU^assembly^_mut-wt_ Calculation with Backrub Sampling

Varying numbers of layers of fibrils protofilaments were prepared for use in in silico estimation of assembly energies. A selection of tau fibril structures was retrieved from the Protein Data Bank (PDB) including PDB IDs: 7p6d, 7p6e, 6vha, 6vh7, 6tjo, 6tjx, 5o3l, 5o3t, 6hre, 7qjw, 6nwp, 6nwq, 6hrf, 7ql4, 5o3o, 5o3t, 7p65, 7p66, 7p67, 7p68, 6gx5, 7p6c, 7p6a, 7p6b, 6qjh, 6qjm, 6qjp and 6qjq. For fibrils with two symmetric protofilaments, a single protofilament was selected for in silico analysis to reduce required computational resources. Using Pymol (version 2.4), fibril PDB structures were created with numbers of layers varying between three and nine using the above mentioned PDB-deposited structures [58]. The deposited and symmetrized assemblies were used to generate assemblies ranging from three to nine layers. Briefly, we used structural alignment to superimpose the top two chains of the deposited fibril with the bottom two chains from a duplicated fibril assembly, preserving the geometry of the assembly while extending the fibril length. Overlapping chains were removed and chains were renamed to produce assemblies of the desired number of layers with chain lettering increasing from the top to the bottom layer. These assemblies were then used as input for the subsequent mutagenesis and minimization using the RosettaScripts interface to Rosetta [59].

Changes in assembly energy were calculated using a method adapted from the Flex ddG protocol described by Barlow *et. al.* [50]. First, the input assembly was placed into a folder along with text files describing the chains of the PDB to mutate to alanine and sets of chains that defined a subunit interface (used for ΔΔG^interface^ calculations). For both the interface and the ΔREU^assembly^_mut-wt_ calculations, all chains were mutated. For the ΔΔG^interface^ calculations on nine-mer assemblies presented, the chains of the center three layers of the nine-mer were used to define the interface. From the input assembly, a set of pairwise atom constraints with a maximum distance of 9 Angstrom were generated with a weight of 1, using the fa_talaris2014 score function. Using this constrained score function, the structure then underwent minimization. After minimization, the residues within 8 angstroms of the mutation site underwent backrub sampling to better capture backbone conformational variation. These sampled structures were either only repacked and minimized, or the alanine mutation was introduced, followed by repacking and minimization. The Rosetta InterfaceDdgMover was used as in the Flex ddG protocol to allow analysis of the ΔΔG^interface^ by defining a fibril interface, giving the lowest energy bound and unbound states for both a wildtype and mutant fibril. For ΔREU^assembly^_mut-wt_ calculations, the bound wild-type and bound mutant structures reported by the interface ddG mover were used for estimating the change in assembly energy due to an alanine substitution. This is repeated for thirty-five independent replicates. The lowest energy bound mutant and bound wild-type structure energies from each replicate were extracted, and the change in energy as calculated by subtracting the wild-type, non-mutagenized assemblies’ energy from the mutant assemblies’ energy. The mean change in energy over the 35 replicates was reported as that residue’s ΔREU^assembly^_mut-wt_. To calculate ΔΔG^interface^ for a residue, the Flex ddG protocol was used as described by Barlow *et al.* [50]. Briefly, the ΔG_wt_ of binding was calculated by subtracting the bound state energy from the unbound state energy for both a wildtype and mutant fibril, and the 1′G_wt_ was subtracted from the ΔG_mut_ to yield the ΔΔG^interface^ for that alanine mutant.

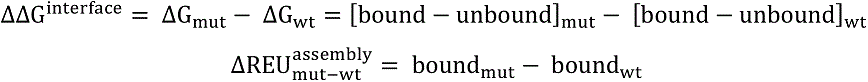

The ΔΔG^interface^ and ΔREU^assembly^_mut-wt_ for a given residue were then averaged over 35 replicates to yield the final values for the residue. This procedure is repeated for every residue in the structure to generate a set of ΔΔG^interface^ and ΔREU^assembly^_mut-wt_ values for all residues in each fibril structure.

### Structure Backbone RMSD analysis

Backbone RMSD analysis of the output structures was carried out by extracting structures used to calculate each replicate’s change in assembly energy (the lowest energy bound mutant and bound wildtype structures for each replicate), using the Flex ddG extract_structures.py script [50]. The extracted structures then had their RMSDs calculated for the PDB chains of interest using the BioPython PDB module to create a list of the backbone atoms (c-α, amide nitrogen, carbonyl carbon, and carbonyl oxygen) for each residue in the chains of interest of both the input structure and the generated structures and using the BioPython PDB.Superimposer module to calculate the RMSD between the two sets of backbone atoms [60]. The resulting RMSDs were then saved by residue for each structure for all thirty-five replicates, allowing by residue and by structure visualization of the structural deformation allowed by the alanine-scanning protocol.

### Per-Residue ΔSASA calculations

Per-residue 1′SASA calculation was performed by utilizing three states for each fibril- the extended monomer (a linear peptide fragment of the fibril generated by using the Pymol ‘fab’ command with the fibril sequence), a monomer in the fibril conformation extracted from the CryoEM fibril structure, and a trimer of the fibril. The SASA of each state was calculated in Rosetta, with the RosettaScripts ‘TotalSasa’ mover with the ‘report_per_residue_sasa’ flag set to true [59]. This generated a per-residue SASA for each residue of the extended monomer, fibril conformation monomer, and for all the residues in the fibril multimer. For the fibril multimer, the SASA of the residues in the central chains was extracted. Then, the per-residue ΔSASA^monomer^_folding_ was calculated by subtracting each residue’s SASA in the folded monomer from the SASA in the extended monomer, and the ΔSASA^in fibril^_folding_ was calculated by subtracting each residue’s SASA in the folded monomer in the center of the fibril from the SASA in the extended monomer.

### Preparation of lysates from CBD patient material

Ten percent weight by volume brain homogenates were made form frontal cortex of CBD cases using Power Gen 125 tissue homogenizer (Fischer Scientific) in 1XTBS buffer supplemented with cOmplete ULTRA protease inhibitor cocktail tablets, EDTA free (Roche). The lysates were sonicated in a water bath (QSonica) for 10 minutes, for 1 minute “on” and 30 seconds “off” intervals, at an amplitude of 65, and centrifuged at 21,000 RCF for 15 minutes at 4°C to remove the debris. The supernatant protein concentration was quantified using a Pierce 660nm Protein Assay Reagent (Thermo Scientific), and subsequently used in seeding assays (see below).

### Construction of WT tauRD (246-408) cell line, seed amplification, and incorporation assay

Lentivirus containing the human WT tauRD (residues 246-408), C-terminally fused to cyan fluorescent protein (CFP), was made in order to constitutively express the WT tauRD-CFP protein in HEK 293-T cells. Lentivirus was made by incubating 7.5uL of TransIT (Mirus) with 142.5uL of Opti-MEM (GIBCO) for 5 minutes. psPAX2 (1200ng), VSV-G (400ng), and WT tauRD-CFP FM5-CMV plasmid (400ng) were mixed into the TransIT/Opti-MEM suspension and incubated for another 25 minutes before addition into a 6 well dish containing HEK 293T cells that were plated with 300,000 cells, 24 hours prior. Cell media (10% FBS, 1% Pen/Strep, 1% GlutaMax in Dulbecco’s modified Eagle’s medium) containing virus was collected after 48 hours, centrifuged at 100 RCF for 5 minutes, and aliquoted before freezing at −80C. HEK 293-T cells were then transduced with the lentivirus and single cell sorted into 96-well plates after expression of protein was confirmed. The cell line was expanded and used for downstream incorporation experiments.

Mutant tauRD (residues 246-408) plasmids, C-terminally fused to mEOS3.2, encoding the top 10 (alanine mutations at amino acid positions 308, 337, 346, 309, 313, 297, 310, 312, 329, and 328) and bottom 10 (alanine mutations at amino acid positions 316, 284, 317, 336, 329, 348, 345, 349, 324, and 322) were synthesized by Twist Biosciences. Virus of each construct was produced by the same lentivirus production protocol outlined above.

In order to amplify the number of aggregates in a population of cells, we treated WT tauRD-CFP cells with CBD brain lysate followed by serial rounds of cell lysis and retreatment of clarified cell lysate onto WT tauRD-CFP cells. Briefly, cells were plated in 96-well plates at 25,000 cells per well in 130 µL of media 12-16 hours before treatment with clarified CBD brain lysate. Initial treatment of WT tauRD-CFP with CBD brain lysate was done by incubating Lipofectamine 2000 (0.75uL per well) with Opti-MEM (19.25 uLper well) for 5 minutes, before adding 5ug of CBD lysate and incubating for another 30 minutes. After treatment, WT tauRD-CFP cells were incubated for 48 hours before cell lysis was performed. Treated cells were harvested with 0.25% trypsin (GIBCO), quenched with cell media and spun down at 100RCF before being resuspend in 1XTBS buffer supplemented with cOmplete ULTRA protease inhibitor cocktail tablets, EDTA free. The resuspended cells were subjected to 3 freeze thaw cycles. The lysates were then sonicated, centrifuged, and quantified using the same protocol used for to prepare the CBD brain lysate. The clarified cell lysate was later used as treatment onto larger well dishes (24 well plates and 6 well plates) until 60-80% of cells contained aggregates. The cells were replated into 96-well plates at 25,000 cells per well in 130 µL of media 12-16 hours before treatment with 10uL tauRD alanine mutants and control lentivirus, in triplicates.

Following 48 hours of lentivirus treatment, the cells were harvested by 0.25% trypsin digestion for 5 min, quenched with cell media, transferred to 96 well U-bottom plates and centrifuged for 5 min at 200 RCF. The cells were then fixed in PBS with 2% paraformaldehyde for 10 minutes, before a final centrifugation step and resuspension in 150uL of PBS. A BD-LSR Fortessa Analyzer instrument was used to perform FRET flow cytometry analysis.

Initial gates were made in order to screen for a population of events that were single cells and double positive for both donor fluorophore (CFP) and acceptor fluorophore (non-photoconverted mEOS3.2). FRET between fluorophores was measured by exciting cells with 405nm violet laser, and emission was collected using a 505nm long pass filter, and 525/50 nm band pass filter.

FCS files were exported from the FACSDiva data collection software and analyzed using FlowJo v10 software (Treestar). Compensation was manually applied to correct for donor bleed-through into the FRET channel guided by a sample with non-aggregated tauRD-mEOS3.2. After selecting for single and double fluorophore positive events, samples were gated on the acceptor intensity such that cells with similar concentrations of tauRD-mEOS3.2 were analyzed to control for the contribution of variable tauRD concentrations to changes in incorporation of alanine mutants. The gating strategy for the FRET experiments is illustrated in Supplementary Fig. 4f.

### Generation of Distance Matrix

To cluster the tau fibril structures, the ΔREU^assembly^_mut-wt_ values for each residue in the fibril were first normalized, and residues not common to all fibrils in the analysis were removed. These normalized ΔREU^assembly^_mut-wt_ values were then used to calculate a distance matrix between each pair of fibril structures using the Scipy library’s ‘spatial.distance_matrix’ method [61].

### Structure Clustering

Agglomerative clustering using Ward’s method and a Euclidean distance metric was performed on the normalized ΔREU^assembly^_mut-wt_ values obtained from the *in silico* alanine scan. Residues not common to all the fibrils being clustered were removed. Using the Scipy library’s ‘cluster.hierarchy.linkage’ method, a linkage matrix describing the clustering of the structures was generated [61]. This linkage matrix was used to generate a dendrogram and plotted alongside the normalized ΔREU^assembly^_mut-wt_ values using the Seaborn library’s clustermap functionality [62, 63].

### Identification of distinguishing fibril features

To identify features distinguishing fibrils, a Random Forest Classification model was generated and interpreted. The normalized ΔREU^assembly^_mut-wt_ values for each of the thirty-five replicates obtained for each residue in the fibril were classified based on the disease of origin (AD, CBD, CTE, PiD, PSP, GGT, GPT). Residues not found in all the structures were removed. Similar residues were grouped together using the Scikit-Learn FeatureAgglomeration functionality to recursively cluster together residues that behaved similarly in the *in silico* alanine scan across the different structures, using Ward’s method with a Euclidean distance metric [64]. The ΔREU^assembly^_mut-wt_ values were in this manner agglomerated to twenty clusters, each valued at the mean of the ΔREU^assembly^_mut-wt_ values of the residues in the cluster.

This clustered ΔREU^assembly^_mut-wt_ data was then split into a test dataset (a random twenty-six of the thirty-five replicates for each residue in the structure) and a train dataset (the remaining nine of the thirty replicates for each residue in the structure). A Random Forest Classifier with 100 classifiers was then trained on the train dataset and classification performance was calculated using the train dataset. This test-train splitting, classifier training, and scoring was repeated 2500 times and the best scoring classifier was taken.

To identify the features being used by the classifier to distinguish the different fibril morphologies, the Python ‘treeinterpreter’ library was used to parse out the contribution of each cluster to the classifier’s prediction of a given fibril type [65]. To do this, the ΔREU^assembly^_mut-wt_ values produced by the *in silico* alanine scan for each fibril (averaged over all thirty-five replicates) was classified by the model, and treeinterpreter was used to extract the contributions of each cluster to correct classification by the Random Forest classification model (i.e., contribution of clusters to classification as AD for AD-PHF/SF fibrils, classification as GGT for GGT_T1/T2/T3 fibrils, etc.).

## Data and code availability

All ThT aggregation data and ΔREU^;<<=16>?^ calculations for tau fibrils are available in https://zenodo.org/record/6407337. All other data are available from the authors upon reasonable request. A protocol capture and code used for this work is made available on https://git.biohpc.swmed.edu/s184069/flex_ddg_ala_scn_runner.

## Acknowledgements

L.A.J. is supported by an Effie Marie Cain Scholarship in Medical Research. C.L.W., M.I.D. and L.A.J. are supported by a Chan Zuckerberg Initiative (CZI) Collaborative Science Award (2018-191983). We thank the members of the Joachimiak lab and Sofia Bali, in particular, for discussions and feedback on the manuscript.

## Author Contributions

V.M., J.V.A. and L.A.J. initiated the project. Peptide aggregation experiments were performed by V.M. TEM of fibrils was performed by B.D.R. V.M. developed the methods for *in silico* minimization of fibril assemblies, estimation of the energy for WT and mutant assemblies, and data analysis. V.M. developed the methods for machine learning classification of features derived from the energies with the help of A.R.V. Validation of the tau alanine mutants in cells was performed by J.V.A. and V.B., using tauopathy patient samples collected and characterized by C.L.W. Finally, V.M., J.V.A., and L.A.J. conceived of and directed the research as well as wrote the manuscript. All authors contributed to the revisions of the manuscript.

## References

1. Eisenberg, D.S. and M.R. Sawaya, Structural Studies of Amyloid Proteins at the Molecular Level. Annual Review of Biochemistry, 2017. 86(1): p. 69–95.

2. Nelson, R., et al., Structure of the cross-β spine of amyloid-like fibrils. Nature, 2005. 435(7043): p. 773-778.

3. Chiti, F. and C.M. Dobson, Protein Misfolding, Amyloid Formation, and Human Disease: A Summary of Progress Over the Last Decade. Annual Review of Biochemistry, 2017. 86(1): p. 27–68.

4. Knowles, T.P.J., M. Vendruscolo, and C.M. Dobson, The amyloid state and its association with protein misfolding diseases. Nature Reviews Molecular Cell Biology, 2014. 15(6): p. 384–396.

5. Willbold, D., et al., Amyloid-type Protein Aggregation and Prion-like Properties of Amyloids. Chemical Reviews, 2021. 121(13): p. 8285–8307.

6. Peng, C., J.Q. Trojanowski, and V.M.Y. Lee, Protein transmission in neurodegenerative disease. Nature Reviews Neurology, 2020. 16(4): p. 199–212.

7. Goedert, M., Alzheimer’s and Parkinson’s diseases: The prion concept in relation to assembled Aβ, tau, and α-synuclein. Science, 2015. 349(6248): p. 1255555.

8. Solforosi, L., et al., A closer look at prion strains. Prion, 2013. 7(2): p. 99–108.

9. Cohen, M., B. Appleby, and J.G. Safar, Distinct prion-like strains of amyloid beta implicated in phenotypic diversity of Alzheimer’s disease. Prion, 2016. 10(1): p. 9–17.

10. Sanders, David W., et al., Distinct Tau Prion Strains Propagate in Cells and Mice and Define Different Tauopathies. Neuron, 2014. 82(6): p. 1271–1288.

11. Shahnawaz, M., et al., Discriminating α-synuclein strains in Parkinson’s disease and multiple system atrophy. Nature, 2020. 578(7794): p. 273-277.

12. Kovacs, G.G., Chapter 25 - Tauopathies, in Handbook of Clinical Neurology, G.G. Kovacs and I. Alafuzoff, Editors. 2018, Elsevier. p. 355–368.

13. Frost, B. and M.I. Diamond, Prion-like mechanisms in neurodegenerative diseases. Nature Reviews Neuroscience, 2010. 11(3): p. 155–159.

14. Kadavath, H., et al., Folding of the Tau Protein on Microtubules. Angew Chem Int Ed Engl, 2015. 54(35): p. 10347–51.

15. Fitzpatrick, A.W.P., et al., Cryo-EM structures of tau filaments from Alzheimer’s disease. Nature, 2017. 547(7662): p. 185-190.

16. Zhang, W., et al., Heparin-induced tau filaments are polymorphic and differ from those in Alzheimer’s and Pick’s diseases. Elife, 2019. 8.

17. Falcon, B., et al., Novel tau filament fold in chronic traumatic encephalopathy encloses hydrophobic molecules. Nature, 2019. 568(7752): p. 420-423.

18. Zhang, W., et al., Novel tau filament fold in corticobasal degeneration. Nature, 2020. 580(7802): p. 283-287.

19. Arakhamia, T., et al., Posttranslational Modifications Mediate the Structural Diversity of Tauopathy Strains. Cell, 2020. 180(4): p. 633–644.e12.

20. Shi, Y., et al., Structure-based classification of tauopathies. Nature, 2021. 598(7880): p. 359-363.

21. Falcon, B., et al., Tau filaments from multiple cases of sporadic and inherited Alzheimer’s disease adopt a common fold. Acta Neuropathol, 2018. 136(5): p. 699–708.

22. Falcon, B., et al., Structures of filaments from Pick’s disease reveal a novel tau protein fold. Nature, 2018. 561(7721): p. 137-140.

23. Makarava, N. and I.V. Baskakov, The same primary structure of the prion protein yields two distinct self-propagating states. The Journal of biological chemistry, 2008. 283(23): p. 15988–15996.

24. Yang, Y., et al., Cryo-EM structures of amyloid-β 42 filaments from human brains. Science, 2022. 375(6577): p. 167-172.

25. Lu, J.-X., et al., Molecular Structure of β-Amyloid Fibrils in Alzheimer’s Disease Brain Tissue. Cell, 2013. 154(6): p. 1257–1268.

26. Xiao, Y., et al., Aβ(1–42) fibril structure illuminates self-recognition and replication of amyloid in Alzheimer’s disease. Nature Structural & Molecular Biology, 2015. 22(6): p. 499–505.

27. Cao, Q., et al., Cryo-EM structure and inhibitor design of human IAPP (amylin) fibrils. Nature Structural & Molecular Biology, 2020. 27(7): p. 653–659.

28. Cao, Q., et al., Cryo-EM structures of hIAPP fibrils seeded by patient-extracted fibrils reveal new polymorphs and conserved fibril cores. Nature Structural & Molecular Biology, 2021. 28(9): p. 724–730.

29. Guerrero-Ferreira, R., et al., Cryo-EM structure of alpha-synuclein fibrils. Elife, 2018. 7.

30. Boyer, D.R., et al., Structures of fibrils formed by α-synuclein hereditary disease mutant H50Q reveal new polymorphs. Nat Struct Mol Biol, 2019. 26(11): p. 1044–1052.

31. Schweighauser, M., et al., Structures of α-synuclein filaments from multiple system atrophy. Nature, 2020. 585(7825): p. 464-469.

32. Boyer, D.R., et al., The α-synuclein hereditary mutation E46K unlocks a more stable, pathogenic fibril structure. Proceedings of the National Academy of Sciences, 2020. 117(7): p. 3592–3602.

33. Li, B., et al., Cryo-EM of full-length α-synuclein reveals fibril polymorphs with a common structural kernel. Nature Communications, 2018. 9(1): p. 3609.

34. Close, W., et al., Physical basis of amyloid fibril polymorphism. Nature Communications, 2018. 9(1): p. 699.

35. Tycko, R., Amyloid Polymorphism: Structural Basis and Neurobiological Relevance. Neuron, 2015. 86(3): p. 632–645.

36. Haj-Yahya, M. and H.A. Lashuel, Protein Semisynthesis Provides Access to Tau Disease-Associated Post-translational Modifications (PTMs) and Paves the Way to Deciphering the Tau PTM Code in Health and Diseased States. Journal of the American Chemical Society, 2018. 140(21): p. 6611–6621.

37. Mirbaha, H., et al., Inert and seed-competent tau monomers suggest structural origins of aggregation. Elife, 2018. 7.

38. Hou, Z., et al., Biophysical properties of a tau seed. Sci Rep, 2021. 11(1): p. 13602.

39. Sharma, A.M., et al., Tau monomer encodes strains. Elife, 2018. 7.

40. Mirbaha, H., et al., Seed-competent tau monomer initiates pathology in PS19 tauopathy mice. bioRxiv, 2022: p. 2022.01.03.474806.

41. Chen, D., et al., Tau local structure shields an amyloid-forming motif and controls aggregation propensity. Nat Commun, 2019. 10(1): p. 2493.

42. von Bergen, M., et al., Mutations of Tau Protein in Frontotemporal Dementia Promote Aggregation of Paired Helical Filaments by Enhancing Local β-Structure*. Journal of Biological Chemistry, 2001. 276(51): p. 48165–48174.

43. Tsolis, A.C., et al., A consensus method for the prediction of ‘aggregation-prone’ peptides in globular proteins. PloS one, 2013. 8(1): p. e54175–e54175.

44. Fernandez-Escamilla, A.-M., et al., Prediction of sequence-dependent and mutational effects on the aggregation of peptides and proteins. Nature Biotechnology, 2004. 22(10): p. 1302–1306.

45. Oliveberg, M., Waltz, an exciting new move in amyloid prediction. Nature Methods, 2010. 7(3): p. 187–188.

46. Goldschmidt, L., et al., Identifying the amylome, proteins capable of forming amyloid-like fibrils. Proceedings of the National Academy of Sciences, 2010. 107(8): p. 3487.

47. Kyte, J. and R.F. Doolittle, A simple method for displaying the hydropathic character of a protein. J Mol Biol, 1982. 157(1): p. 105–32.

48. Biancalana, M. and S. Koide, Molecular mechanism of Thioflavin-T binding to amyloid fibrils. Biochimica et biophysica acta, 2010. 1804(7): p. 1405–1412.

49. Lövestam, S., et al., Assembly of recombinant tau into filaments identical to those of Alzheimer’s disease and chronic traumatic encephalopathy. bioRxiv, 2021: p. 2021.12.16.472950.

50. Barlow, K.A., et al., Flex ddG: Rosetta Ensemble-Based Estimation of Changes in Protein-Protein Binding Affinity upon Mutation. The journal of physical chemistry. B, 2018. 122(21): p. 5389–5399.

51. Smith, C.A. and T. Kortemme, Backrub-like backbone simulation recapitulates natural protein conformational variability and improves mutant side-chain prediction. Journal of molecular biology, 2008. 380(4): p. 742–756.

52. Nazarov, S., et al., The structural basis of huntingtin (Htt) fibril polymorphism, revealed by cryo-EM of exon 1 Htt fibrils. bioRxiv, 2021: p. 2021.09.23.461534.

53. Hervas, R., et al., Cryo-EM structure of a neuronal functional amyloid implicated in memory persistence in Drosophila. Science, 2020. 367(6483): p. 1230-1234.

54. Abskharon, R., et al., Cryo-EM structure of RNA-induced tau fibrils reveals a small C-terminal core that may nucleate fibril formation. bioRxiv, 2022: p. 2022.01.28.478258.

55. Holmes, B.B., et al., Proteopathic tau seeding predicts tauopathy in vivo. Proceedings of the National Academy of Sciences, 2014. 111(41): p. E4376–E4385.

56. Sanders, D.W., et al., Distinct tau prion strains propagate in cells and mice and define different tauopathies. Neuron, 2014. 82(6): p. 1271–88.

57. Khan, T., et al., Quantifying Nucleation In Vivo Reveals the Physical Basis of Prion-like Phase Behavior. Molecular cell, 2019. 73(4): p. 857–857.

58. Schrodinger, LLC, The PyMOL Molecular Graphics System, Version 2.4/2.5. 2015.

59. Fleishman, S.J., et al., RosettaScripts: A Scripting Language Interface to the Rosetta Macromolecular Modeling Suite. PLOS ONE, 2011. 6(6): p. e20161.

60. Cock, P.J.A., et al., Biopython: freely available Python tools for computational molecular biology and bioinformatics. Bioinformatics, 2009. 25(11): p. 1422–1423.

61. Virtanen, P., et al., SciPy 1.0: fundamental algorithms for scientific computing in Python. Nat Methods, 2020. 17(3): p. 261–272.

62. Hunter, J.D., Matplotlib: A 2D Graphics Environment. Computing in Science & Engineering, 2007. 9(3): p. 90–95.

63. Waskom, M.L., seaborn: statistical data visualization. Journal of Open Source Software, 2021. 6(60): p. 3021.

64. Pedregosa, F., et al., Scikit-learn: Machine Learning in Python. Journal of Machine Learning Research, 2011. 12(85): p. 2825–2830.

65. Saabas, A. treeinterpreter. 2021 [cited 2022 January 12]; Available from: https://github.com/andosa/treeinterpreter.

66. Holehouse, A.S., et al., CIDER: Resources to Analyze Sequence-Ensemble Relationships of Intrinsically Disordered Proteins. Biophys J, 2017. 112(1): p. 16–21.

67. Mutations | MAPT. [cited 2021 November 12]; Available from: https://www.alzforum.org/mutations/mapt.

