## Supplementary Information for "Network of hotspot interactions cluster tau amyloid folds"

### Supplementary Figures and Legends

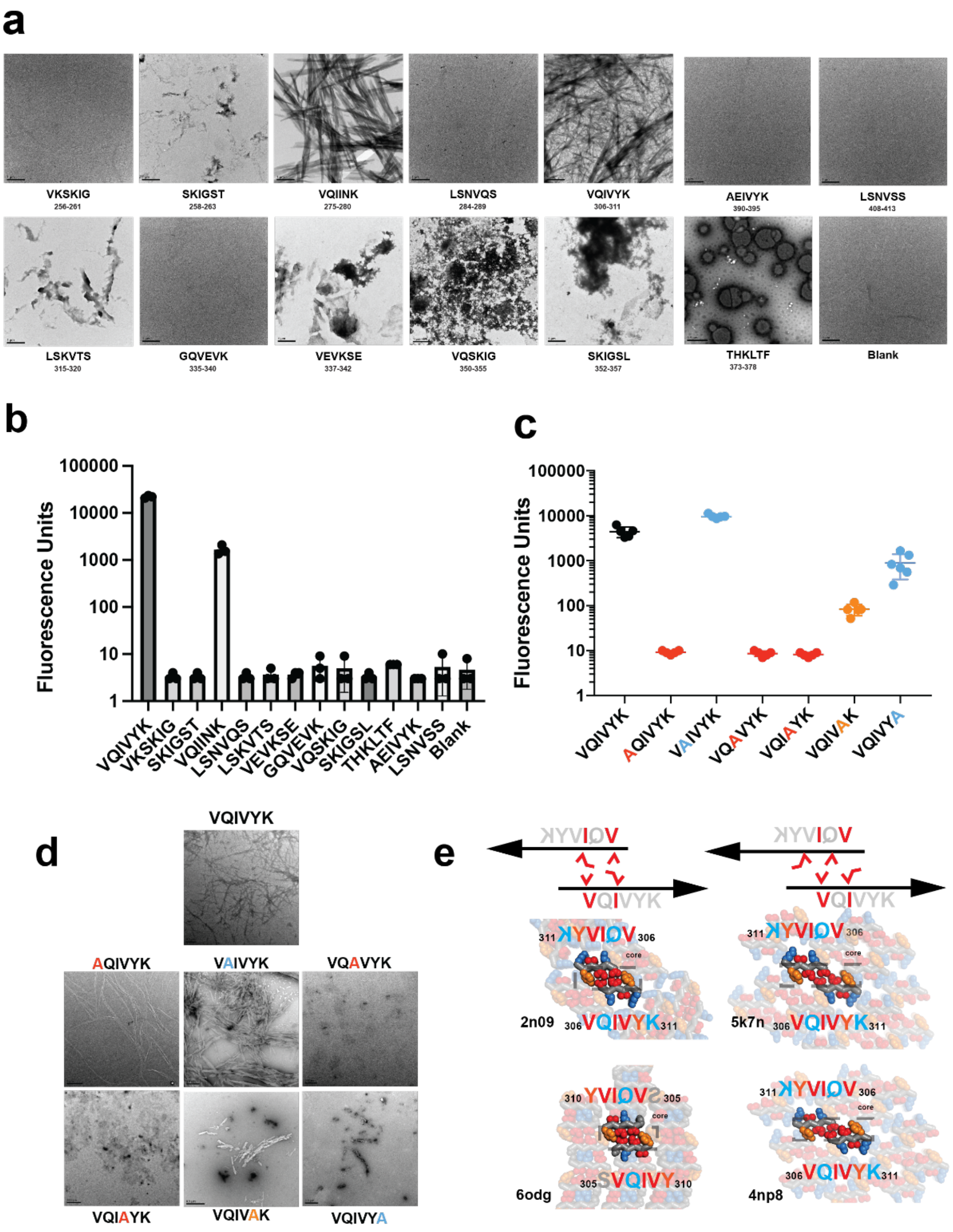


**Supplementary Figure 1. Amyloid properties that determine assembly. (a)** Representative TEM images of amyloid motifs from tau predicted by ZipperDB. Images were acquired from ThT fluorescence aggregation endpoints from **(b)**. Scale bar is 1 μm. **(b)** ThT fluorescence end points for predicted amyloid motifs from tau. Aggregation experiments were performed in triplicate and shown as averages with standard deviation. **(c)** ThT fluorescence aggregation end points for $^{306}$VQIVYK$^{311}$ and alanine mutants at each position. Aggregation experiments were performed as five replicates and are shown as averages with standard deviation. Peptides that yielded high ThT are shown in blue, peptides with no ThT are colored in red and an intermediate mutant is colored in light blue. **(d)** TEM images of VQIVYK and its alanine mutants. Scale bar is 1um. **(e)** Mapping aggregation properties from the alanine mutants onto four available structures of VQIVYK (A-C; PDB IDs: 2n09, 5k7n and 4np4) or SVQIVY (D; PDB id 6odg). Symmetry related lattice of the peptide is shown in spacefill representation and the residues in the core “dimer” are stabilized key residues important for aggregation. The residues are colored by their aggregation signal from **(c)**. The structures are summarized by the relative register of the monomers in the core dimer, in register (left) or off set (right).

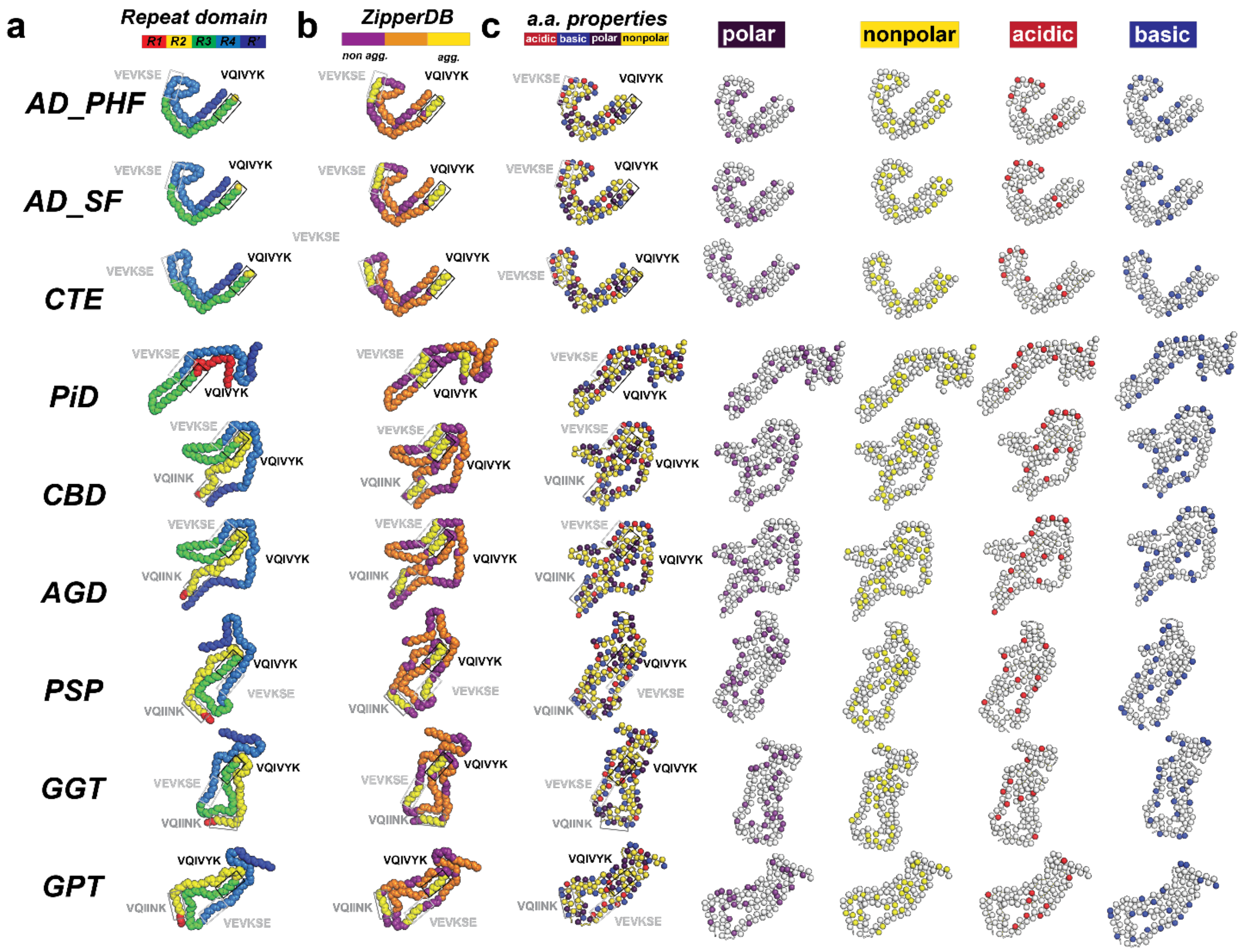


**Supplementary Figure 2. Features of the tauopathy fibril structures. (a)** Fibril structures are colored by repeat domain in red, yellow, green, blue and dark blue for repeat domains R1, R2, R3, R4, and R' respectively. Structures are shown in spacefill representation for the backbone. **(b)** Fibrils are colored by aggregation propensity of the different motifs colored in yellow, orange and magenta for high, medium, and low aggregation propensity. Structures are shown in spacefill for the backbone. **(c)** Fibrils are colored by amino acid type, nonpolar, polar, acidic, and basic are colored yellow, purple, red and blue respectively. Structures are shown using a ribbon representation and the c-β atom is shown in spheres. The $^{306}$VQIVYK$^{311}$, $^{275}$VQIINK$^{280}$, and $^{337}$VEVKSE$^{342}$ motifs are highlighted in a box in each structure. PDB ids: 5o3l, 5o3t 6gx5, 6nwp, 6tjo, 7p6d, 7p65, 7p66 and 7p6a.

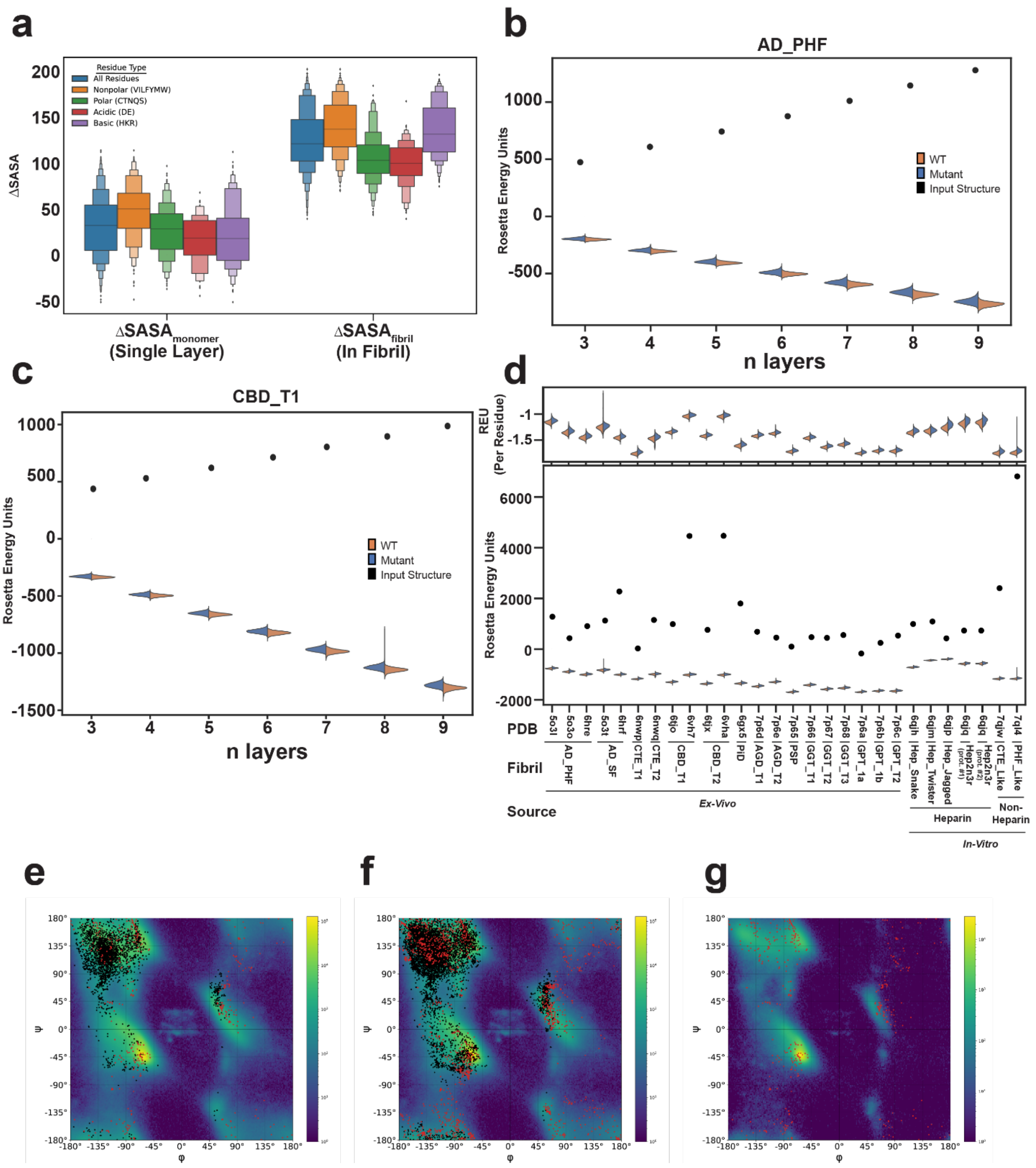


**Supplementary Figure 3. Evaluating stability of fibril assemblies. (a)** Quantification of the change in Solvent Accessible Surface Area (SASA) from the unfolded monomer to the folded monomer ($\Delta SASA^{\mathrm{monomer}}_{\mathrm{folding}}$) (left panel) or to the folded monomer in the fibril ($\Delta SASA^{\mathrm{in\ fibril}}_{\mathrm{folding}}$) (right panel) calculated for all, nonpolar, polar, basic, and acidic residues colored in blue, orange, green, red, and purple. Data is shown as a boxen or 'letter-value' plot for ΔSASA values for nine tauopathy protofilament structures. Evaluation of total energies (REU) for AD-PHFs (PDB id: 5o3l) **(b)** and CBD (PDB id: 6tjo) **(c)** using different numbers of layers from three to nine for all structures produced during the alanine scan. Distribution of energies for WT and alanine mutants are shown in orange and blue, respectively. Energies for native starting structures are shown as black points. **(d)** Total energy distributions for minimized wild-type and alanine mutant nine-mer (bottom) using our protocol for all available tauopathy and four heparin tau fibrils, along with the energies of the input, un-minimized structures (points). The same distributions are shown normalized to the number of residues in the repeating monomer unit (top). The plot is colored as in **(b, c)**. PDB IDs: 7p6d, 7p6e, 6vha, 6vh7, 6tjo, 6tjx, 5o3l, 6hre, 7qjw, 6nwp, 6nwq, 6hrf, 7ql4, 5o3o, 5o3t, 7p65, 7p66, 7p67, 7p68, 6gx5, 7p6c, 7p6a, 7p6b. **(e)** Overlay of φ/ψ torsional distributions for X-ray structures of globular proteins with resolution between 1.5 Å and 2.5 Å (110,677 structures) to φ/ψ torsional angles for all residues (black) and glycine residues (red) in tau fibril structures (35 structures) or **(f)** all fibril structures (96 structures) determined using helical reconstruction using RELION. **(g)** Overlay of alanine φ-ψ torsional distributions (background) for X-ray structures of globular proteins with resolution between 1.5 Å and 2.5 Å with glycine φ/ψ torsional angles (red) in tau fibrils determined using helical reconstruction in RELION. ΔSASA distributions are shown as letter-value plots with the center 2 boxes showing 50% of the data with each smaller box contain half of the remaining data. REU energy distributions are shown across 35 replicates at each position in each fibril and plotted as violin plots.

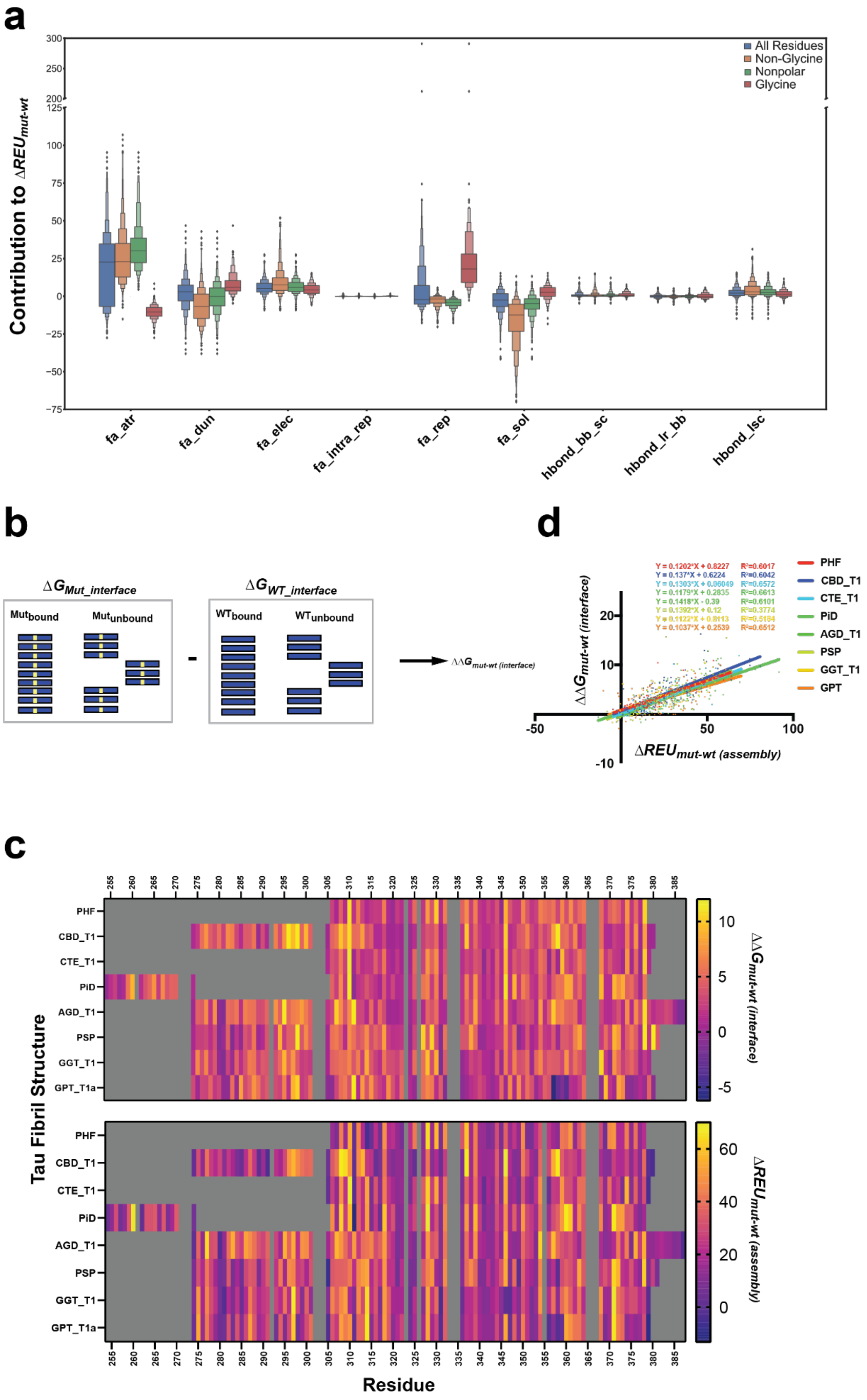

a
Contribution to ΔREU_mut-wt
All Residues
Non-Glycine
Nonpolar
Glycine
300
250
200
125
100
75
50
25
0
-25
-50
-75
fa_atr
fa_dun
fa_elec
fa_intra_rep
fa_rep
fa_sol
hbond_bb_sc
hbond_lr_bb
hbond_lsc
b
ΔG_Mut_interface
Mut_bound
Mut_unbound
ΔG_WT_interface
WT_bound
WT_unbound
ΔΔG_mut-wt (interface)
d
Y = 0.1202*X + 0.8227 R²=0.6017
Y = 0.137*X + 0.6224 R²=0.6042
Y = 0.1303*X + 0.06049 R²=0.6572
Y = 0.1179*X + 0.2835 R²=0.6613
Y = 0.1418*X - 0.39 R²=0.6101
Y = 0.1392*X + 0.12 R²=0.3774
Y = 0.1122*X + 0.8113 R²=0.5184
Y = 0.1037*X + 0.2539 R²=0.6512
PHF
CBD_T1
CTE_T1
PiD
AGD_T1
PSP
GGT_T1
GPT
ΔΔG_mut-wt (interface)
ΔREU_mut-wt (assembly)
20
10
-10
-50
50
100
c
Tau Fibril Structure
PHF
CBD_T1
CTE_T1
PiD
AGD_T1
PSP
GGT_T1
GPT_T1a
ΔΔG_mut-wt (interface)
10
5
0
-5
ΔREU_mut-wt (assembly)
60
40
20
0
Residue
255 260 265 270 275 280 285 290 295 300 305 310 315 320 325 330 335 340 345 350 355 360 365 370 375 380 385

**Supplementary Figure 4. Evaluation of per residue energetics in fibrils. (a)** Breakdown of energy terms from Rosetta energy calculations evaluating their overall contribution to the assembly stability across nine different tauopathy fibril structures. Distributions for all, non-glycine, nonpolar and glycine are colored in blue, orange, green and red, respectively. Energy term distributions are shown across 35 replicates across positions clustered by amino acid properties. Energy distributions are shown as letter-value plots with the center 2 boxes showing 50% of the data with each smaller box contain half of the remaining data. **(b)** Schematic for the Flex ddG-based protocol for determining the contribution of residues to the energetics of inter-layer interfaces in protein fibrils. **(c)** Correlations between the *in silico* estimation of assembly stability upon mutation (x-axis) and the *in silico* "interface" inter-layer contribution of the assembly stability upon mutation (y-axis) for tauopathy fibrils. PDB ids: 5o3l, 6gx5, 6nwp, 6tjo, 7p6d, 7p65, 7p66 and 7p6a. **(d)** Heatmap comparison of the energetic change in response to substitution to alanine of the interface within a tau fibril (top) or of the total structural energy (bottom) as predicted by Rosetta. Scale is colored in the plasma color scheme from yellow (most important) to purple (least important).

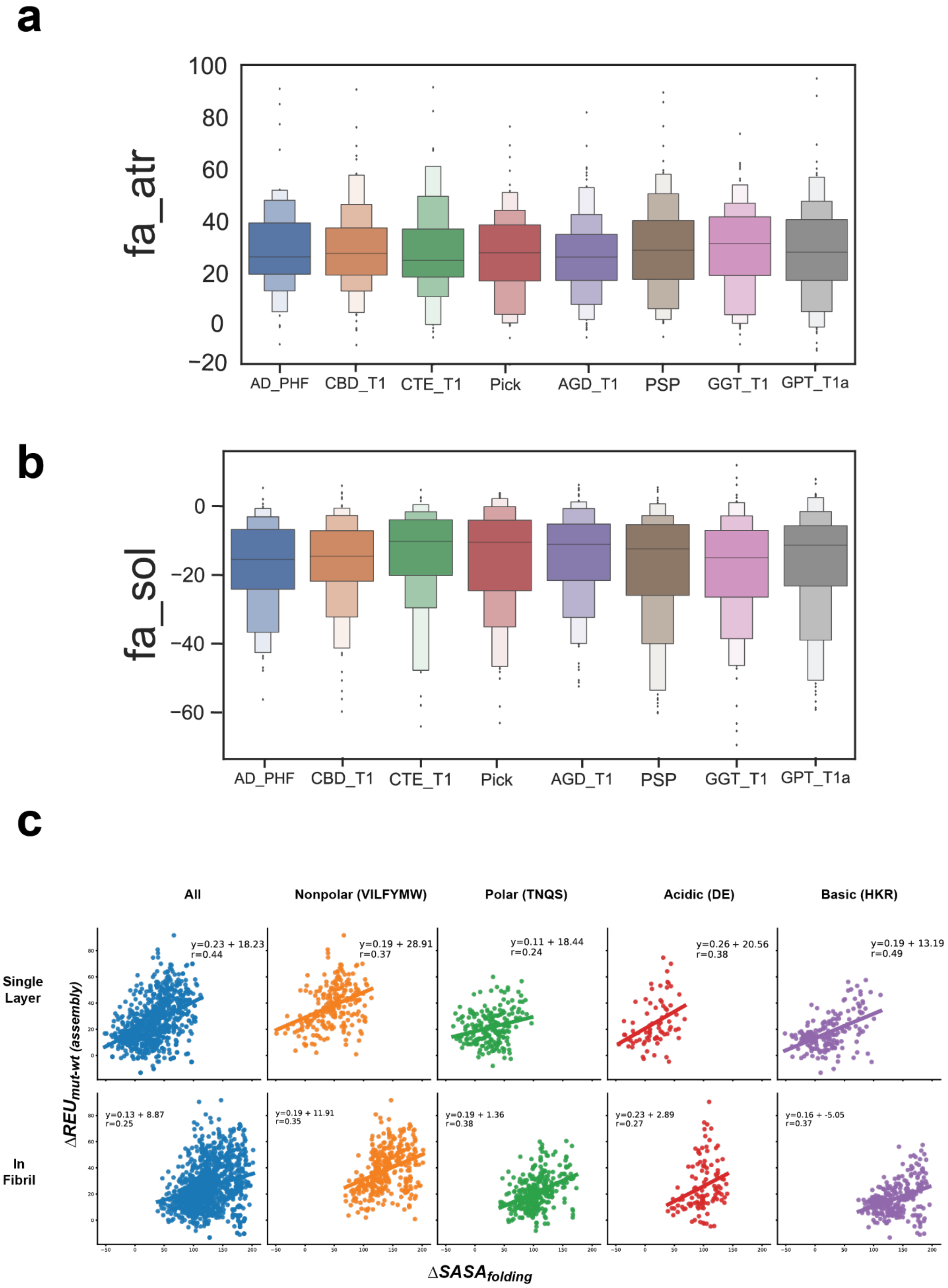

a
fa_atr
100
80
60
40
20
0
−20
AD_PHF
CBD_T1
CTE_T1
Pick
AGD_T1
PSP
GGT_T1
GPT_T1a
b
fa_sol
0
−20
−40
−60
AD_PHF
CBD_T1
CTE_T1
Pick
AGD_T1
PSP
GGT_T1
GPT_T1a
c
All
Nonpolar (VILFYMW)
Polar (TNQS)
Acidic (DE)
Basic (HKR)
Single Layer
In Fibril
y=0.23 + 18.23
r=0.44
y=0.19 + 28.91
r=0.37
y=0.11 + 18.44
r=0.24
y=0.26 + 20.56
r=0.38
y=0.19 + 13.19
r=0.49
y=0.13 + 8.87
r=0.25
y=0.19 + 11.91
r=0.35
y=0.19 + 1.36
r=0.38
y=0.23 + 2.89
r=0.27
y=0.16 + -5.05
r=0.37
ΔREU mut-wt (assembly)
ΔSASA folding

**Supplementary Figure 5. Energy contributions and solvent accessibility to stabilizing fibril conformations.** Comparison of the contribution of the full atom Lennard-Jones attractive potential energy (fa_atr) **(a)** and the full atom Lazaridis-Karplus solvation energy (fa_sol) (**b**) as calculated by Rosetta to the *in silico* predicted energy for the assembly of selected tauopathy fibril structures. Contributions are plotted in Rosetta Energy Units and are shown as letter-value plots with the center 2 boxes showing 50% of the data with each smaller box contain half of the remaining data. **(c)** Correlations between the ΔSASA of folding and the *in silico* change in assembly energy upon mutation (y-axis) for all, nonpolar, polar, acidic, and basic residues, either in the context of a single monomer folded into its fibril conformation without (top) or within (bottom) the context of the fibril. PDB ids: 5o3l, 6gx5, 6nwp, 6tjo, 7p6d, 7p65, 7p66 and 7p6a.

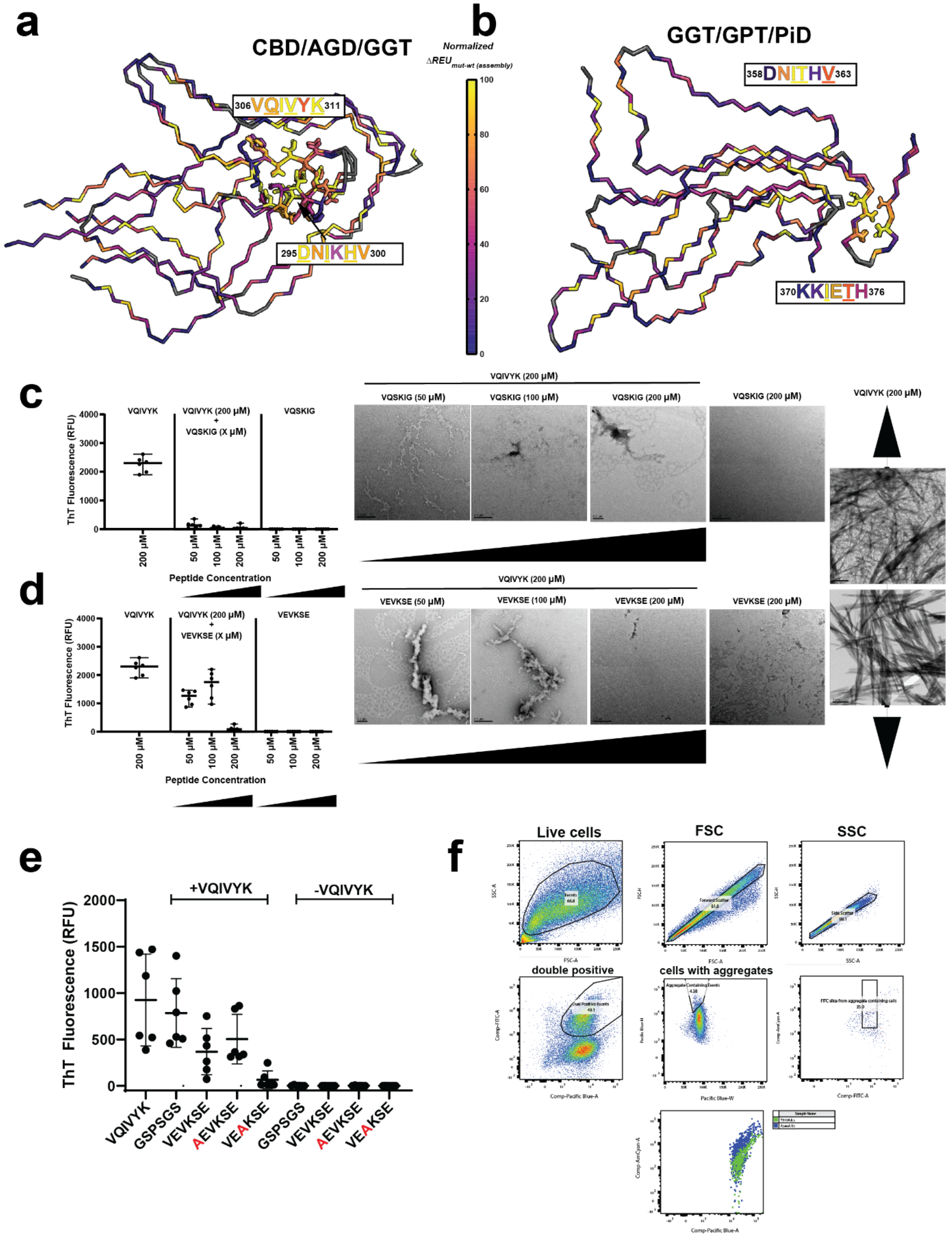


**Supplementary Figure 6. Competing peptides interfere with VQIVYK assembly. (a)** Alignment of CBD/AGD/GGT tauopathy fibril monomer conformations on the $^{306}$VQIVYK$^{311}$ amyloid-forming motif **(b)** and GGT/GPT/Pick's disease fibril monomer conformations on $^{358}$DNITHV$^{363}$. Residues are colored in the plasma color scheme from yellow (important) to purple (unimportant) by their effect on *in silico* assembly stability when mutated to alanine. Structures are shown as ribbons and key residues are shown in stick representation. Endpoint Thioflavin T fluorescence values for 200 μM $^{306}$VQIVYK$^{311}$ peptide aggregation (left), 200 μM $^{306}$VQIVYK$^{311}$ with 50, 100 or 200 μM competitor (middle) and 50, 200 and 200 μM competitor alone (right). Competitor peptides $^{350}$VQSKIG$^{355}$ (**c**) and $^{337}$VEVKSE$^{342}$ (**d**) were co-aggregated with $^{306}$VQIVYK$^{311}$. Experiments were performed as six technical replicates and are reported as averages with standard deviation. TEM images of each endpoint sample are shown on the right. Scale bars are shown as 0.2 μm. (**e**) Aggregation of 50 μM $^{337}$VEVKSE$^{342}$, VEVKSE alanine mutants (AEVKSE and VEAKSE) and GSPSGS control peptide with and without the presence of 200 μM $^{306}$VQIVYK$^{311}$ peptide. Experiment was performed as six technical replicates. Bars report average and standard deviation.(**f**) Gating strategy for flow cytometry readout of in-cell incorporation assay on tau alanine mutants.

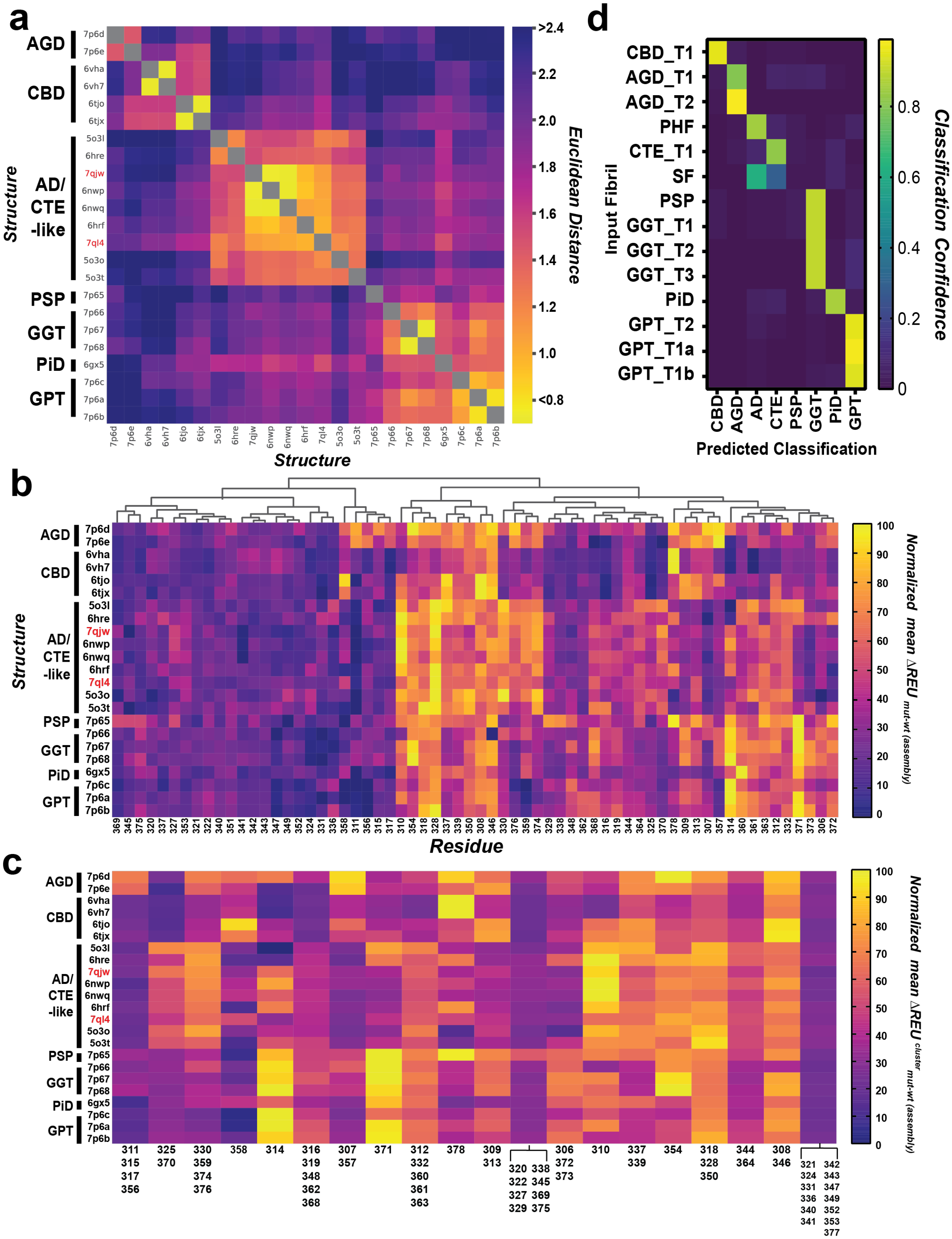


**Supplementary Figure 7. ML-based classification of features based on residue energetics. (a**) Similarity matrix of *ex vivo* fibrils based on *in silico*

$\Delta\mathrm{REU}_{\mathrm{mut-wt}}^{\mathrm{assembly}}$ measurements. Cells colored with the plasma color scheme by Euclidean distance between vectors of fibril pairs' $\Delta\mathrm{REU}_{\mathrm{mut-wt}}^{\mathrm{assembly}}$ measurements from yellow (<0.8 REU, most similar) to blue (>2.4, most dissimilar). **(b)** Dendrogram generated by hierarchical clustering via Ward's method of fibril residues, displayed alongside heatmap of *in silico* $\Delta\mathrm{REU}_{\mathrm{mut-wt}}^{\mathrm{assembly}}$. Dendrogram groups residues that covary by *in silico* $\Delta\mathrm{REU}_{\mathrm{wt-mut}}^{\mathrm{assembly}}$ across structures together. Residues are colored in the heatmap with the plasma color scheme from yellow (highest *in silico* $\Delta\mathrm{REU}_{\mathrm{mut-wt}}^{\mathrm{assembly}}$) to purple (lowest *in silico* $\Delta\mathrm{REU}_{\mathrm{mut-wt}}^{\mathrm{assembly}}$). **(c)** Heatmap of residue clusters generated via feature agglomeration, recursively combining the most similar residues/residue clusters together. Clusters are colored by the arithmetic mean value of residues comprising the cluster from yellow (highest mean *in silico* $\Delta\mathrm{REU}_{\mathrm{mut-wt}}^{\mathrm{assembly}}$) to purple (lowest mean *in silico* $\Delta\mathrm{REU}_{\mathrm{mut-wt}}^{\mathrm{assembly}}$). **(d)** Classification confidence matrix of the Random Forest Classifier when predicting classifications using the clustered mean *in silico* $\Delta\mathrm{REU}_{\mathrm{mut-wt}}^{\mathrm{assembly}}$ values derived from the *in silico* alanine scan. When given an input fibril structure's clustered mean *in silico* $\Delta\mathrm{REU}_{\mathrm{mut-wt}}^{\mathrm{assembly}}$ values, the squares represent the class probabilities the random forest classifier assigns to the input, colored with the plasma color scheme from yellow (1.0 probability, the classifier model is highly certain of a class assignment to the input) to purple (0 probability, the classifier model does not predict the input to belong to a given class). PDB IDs: 7p6d, 7p6e, 6vha, 6vh7, 6tjo, 6tjx, 5o3l, 6hre, 7qjw, 6nwp, 6nwq, 6hrf, 7ql4, 5o3o, 5o3t, 7p65, 7p66, 7p67, 7p68, 6gx5, 7p6c, 7p6a, 7p6b.
